# Modelling the mechanical cross-talk between cells and fibrous extracellular matrix using hybrid cellular Potts and molecular dynamics methods

**DOI:** 10.1101/2022.06.10.495667

**Authors:** Erika Tsingos, Bente Hilde Bakker, Koen A.E. Keijzer, Hermen Jan Hupkes, Roeland M.H. Merks

**Author notes:** these authors contributed equally to this work (Erika Tsingos, Bente Hilde Bakker).

## Abstract

The mechanical interaction between cells and the extracellular matrix (ECM) is fundamental to coordinate collective cell behavior in multicellular tissues. Relating individual cell-level mechanics to tissue-scale collective behavior is an outstanding challenge which cell-based models such as the cellular Potts model (CPM) are well-positioned to address. These models generally represent the ECM with mean-field approaches, which assume substrate homogeneity. This assumption breaks down with fibrous ECM, which has non-trivial structure and mechanics. Here, we extend the CPM with a bead-spring chain model of ECM fiber networks modelled using molecular dynamics. We model contractile cells pulling with discrete focal adhesion-like sites on the ECM fiber network, and demonstrate agreement with experimental spatiotemporal fiber densification and displacement. We show that contractile cell forces propagate over multiple cell radii scaling with power law exponent of ≈ −0.5 typical of viscoelastic ECM. Further, we use in silico atomic force microscopy to measure local cell-induced network stiffening consistent with experiments. Our model lays the foundation to investigate how local and long-ranged cell-ECM mechanobiology contributes to multicellular morphogenesis.

## Introduction

Cells and tissues are in a constant reciprocal mechanical cross-talk with the extracellular matrix (ECM), whose major structural and load-bearing components are fibrous proteins or proteoglycans such as collagens and fibronectins that assemble into complex networks [Walma and Yamada, 2020]. The density of fibers, their alignment, and density of fiber crosslinking affect material stiffness and create non-linear mechanical properties [Chaudhuri et al., 2020, Notbohm et al., 2015]. Cells sculpt ECM fibers via traction forces, and in turn fiber network mechanics impinge on cells’ cytoskeleton, altering cell shape and fate [Dzamba and DeSimone, 2018, Van Helvert et al., 2018]. The fiber network enables transmission of forces over distances of several cells [Wang and Xu, 2020], coordinating morphogenesis at the cell and tissue scale [Korff and Augustin, 1999, Naganathan et al., 2022, Palmquist et al., 2022].

Several sophisticated mechanical models of ECM fiber networks have been developed (reviewed by Alisafaei et al. [2021], Sree and Tepole [2020], Wang and Xu [2020]). By design, these approaches focus on the ECM’s material properties, and are thus limited in two respects. First, they oversimplify biological behavior of cells that are embedded into and interact with the fiber network [Sree and Tepole, 2020]. Second, they struggle with the mesoscale: They can only capture cell-ECM mechanobiology when cell density is very low [Eichinger et al., 2021, Guo et al., 2022], or, conversely, when cell density is very high such that multicellular collectives can be approximated by a continuum [Guo et al., 2022]. Cell-based models are naturally well-adapted to address both of these limitations. They are a versatile family of models for discrete, mechanically interacting cells with applications ranging from cancer biology [Metzcar et al., 2019], evolutionary biology [Colizzi et al., 2020], to developmental biology and morphogenesis [Rens et al., 2020]. Generally, these models treat cell-ECM mechanobiology implicitly by coarse-graining ECM fibers as a homogeneous material exerting uniform friction [Daub and Merks, 2013, Drasdo and Hoehme, 2012], or explicitly with a finite element model [Rens and Merks, 2017, 2020, Scott et al., 2020, Van Oers et al., 2014]. However, the characteristic length scale of ECM fibers is large compared to the cell size; thus, coarse-graining fails to capture some material properties, and a discrete representation is needed [Davidson et al., 2019, Guo et al., 2022, Wang and Xu, 2020]. Several cell-based models represent the ECM discretely, e.g. with collections of point particles [Buske et al., 2012, Dunn et al., 2012, Marin-Riera et al., 2016, Nelemans et al., 2020, Sütterlin et al., 2017] or thin rigid cylinders [Bauer et al., 2007, Macnamara et al., 2020, Peurichard et al., 2017, Schlüter et al., 2012], but without mechanically linking these into fiber networks. A notable exception is the model of Reinhardt and Gooch [2014, 2018], because it represents ECM fibers as serial collections of linear springs. Their model qualitatively reproduces cell-matrix remodelling and collagen bulk mechanics [Reinhardt and Gooch, 2014, 2018]. Thus, coupling cell-based models with spring-based network models of ECM fibers is a promising avenue to investigate how collective multicellular behavior at the tissue scale emerges from cell-ECM mechanobiology at the cell scale.

Among cell-based models, the cellular Potts model (CPM) stands out for its flexible and biophysically realistic treatment of cell shape [Albert and Schwarz, 2014, Hirashima et al., 2017]. Originally developed to test the differential adhesion hypothesis in developmental biology, the CPM traces back to the large q-Potts model in statistical mechanics, and represents cells as a collection of elements on a regular grid [Glazier and Graner, 1993, Graner and Glazier, 1992]. The model’s dynamics are governed by a Hamiltonian function, an expression for the system’s energy that encapsulates all processes that are relevant for the system. In its most common formulation, the Hamiltonian represents the balance of forces for an overdamped Newtonian system; evaluating energy gradients associated with cell shape changes is equivalent to solving the force balance [Hirashima et al., 2017, Rens and Edelstein-Keshet, 2019]. Additional terms inspired by biophysical observations, such as cell surface tension, enable the CPM to accurately predict cell spreading on adhesive substrates, focal adhesion formation, and the stresses generated by cell forces on the ECM [Albert and Schwarz, 2014, Rens and Merks, 2017, 2020, Van Oers et al., 2014]. The framework is amenable to extensions, for example to include force generation and cell shape changes induced by the actin cytoskeleton dynamics [Marée et al., 2006, Niculescu et al., 2015, Schakenraad et al., 2022, Van Steijn et al., 2022]. These features have made the CPM a popular tool to study the emergence of biophysical properties at the cell and tissue scale from simple (sub-)cellular rules [Albert and Schwarz, 2014, Rens and Merks, 2020, Wortel et al., 2021].

Here, we introduce a CPM hybridized with a discrete mechanical model for ECM fiber networks in continuous space. Modelling cells with the CPM allows to benefit from the vast prior biophysical studies using this model. On the other hand, thin sub-micrometer ECM fibers are best represented in continuous space to avoid cumbersome computational checks to keep fibers connected on a discrete lattice. Our model was implemented by interfacing the CPM software library Tissue Simulation Toolkit [Daub and Merks, 2014] with the high-performance molecular mechanics framework HOOMD-blue [Anderson et al., 2008, 2020]. We demonstrate the model’s capabilities by replicating a common experimental setup involving a contractile cell straining a randomly-oriented fiber network. We quantitatively compare our results to published experimental data while varying parameters. We show that ECM parameters for fiber density and crosslinking, as well as cell parameters for density of cell-ECM adhesions and the cell’s intrinsic ability to cluster adhesions determine the extent of cell contraction and force generation, and the transmission of long-ranged forces through the network. Looking forward, our framework can be calibrated to the network structure and mechanics of mixtures of ECM fiber types found *in vivo*, and can incorporate existing CPMs such as the dynamic force-dependent focal adhesion model in Rens and Merks [2020] to scale to multicellular morphogenesis.

## Results

### A cellular Potts model interfaced with discrete ECM fibers

In the following, we briefly delineate our modelling choices and implementation. For a more detailed technical description, see the Methods section.

To model cells, we adapted the 2D CPM [Glazier and Graner, 1993, Graner and Glazier, 1992] as implemented in the Tissue Simulation Toolkit [Daub and Merks, 2013, 2014, Merks and Glazier, 2005]. In the CPM, each cell is represented as a collection of (usually connected) sites in a regular, usually square lattice (Fig. 1 A); any change in the spatial configuration of cell sites is assumed to require physical work. This change in work is described by a Hamiltonian energy function *H* (Equation 13, Methods section), which generally contains terms for the energy of cell-cell adhesion, and deviation from a preferred target area [Daub and Merks, 2013]. We introduced additional terms to this “classical” CPM Hamiltonian to describe the work involved in mechanical cell-ECM interactions. The temporal dynamics of the CPM were evaluated based on energy minimization of the Hamiltonian using Metropolis dynamics [Glazier and Graner, 1993, Graner and Glazier, 1992].

**Figure 1:**
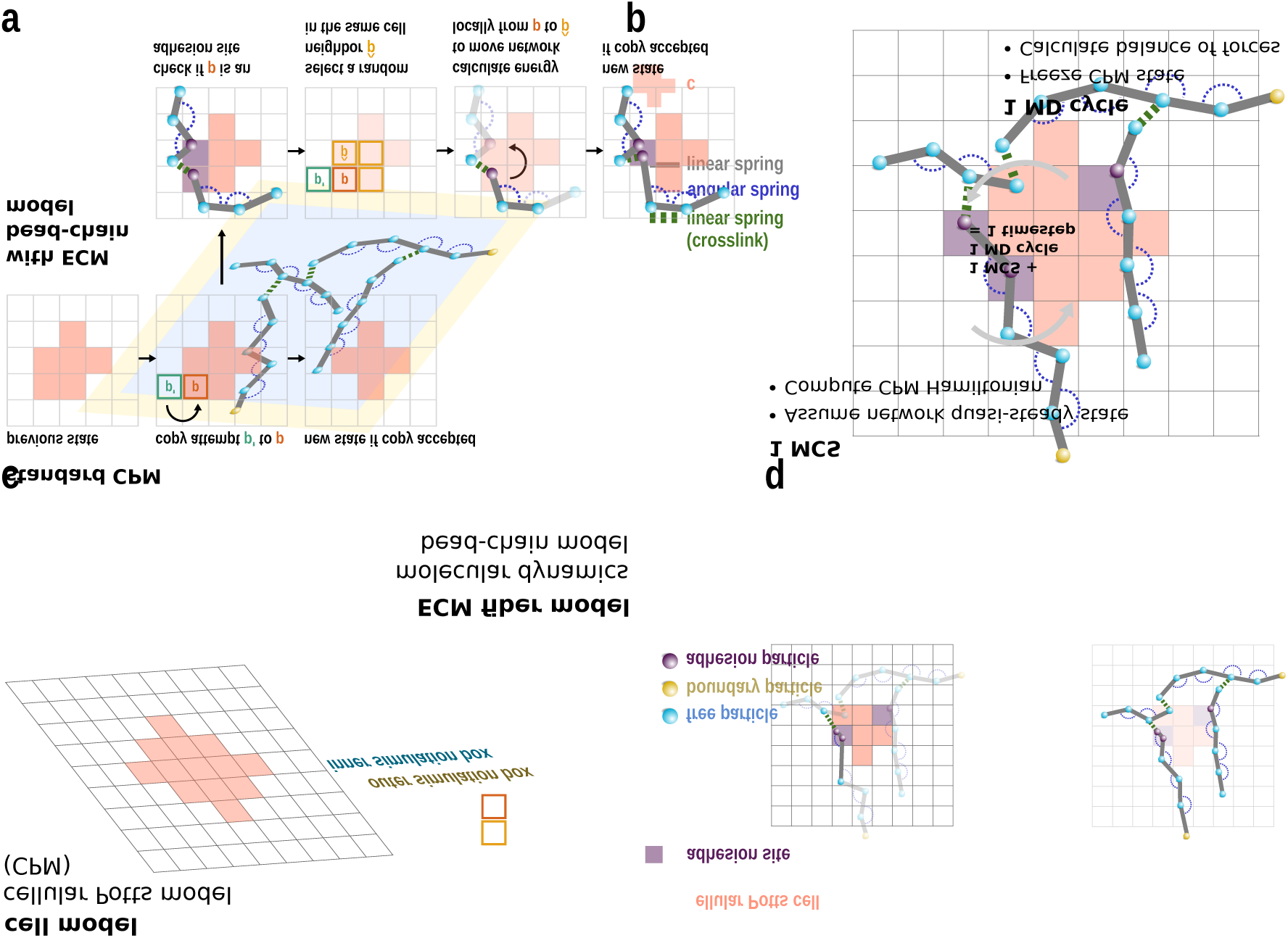
Model overview. **a**. We combine a cellular Potts model (CPM) of cells with a particle-based model of the extracellular matrix (ECM) modelled as a bead-chain polymer network. **b**. Adhesions are the mechanical interface between the two model layers. Adhesions consist of an adhesion site in the CPM layer, and at least one adhesion-associated particle. **c**. CPM lattice sites that are not associated to a cell-ECM adhesion undergo copy attempts according to standard CPM rules (top row). When the lattice site *p* coincided with an adhesion site, any associated adhesion particles were moved to an adjacent site belonging to the same cell 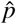. If 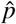already contained adhesion particles, a clustering energy was calculated according to Equation 12. If no valid site 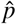 existed, then the adhesion particle was released and became a free particle. **d**. During simulation, we solve the CPM for one Monte Carlo step (MCS) with the assumption that the network has relaxed to a quasi-steady state. After each MCS, we integrate the ECM network model for one molecular dynamics (MD) cycle while keeping the CPM state frozen.

Fiber-forming ECM proteins such as collagen and fibrin self-assemble from nanometer-scale components (fibrils) into fibers with micrometer-scale length but sub-micrometer diameter [Kreger et al., 2010, Leonidakis et al., 2017]. We reasoned that modelling the dynamics of ECM fibers entirely within the CPM (e.g. by introducing dynamics into previous, static representations of ECM fibers as a special type of cell, as in Bauer et al. [2007] or Scianna et al. [2013]) would require excessively reducing the lattice size and introducing computationally expensive checks to ensure fibers retained their connectivity. We therefore opted to model ECM fibers in a separate model interfaced with the CPM (Fig. 1 A). The mechanical behavior of individual collagen fibrils at small strains can be reasonably approximated with a linear elastic spring [Sasaki and Odajima, 1996, Svensson et al., 2010]. Indeed, mechanical models abstracting collagen fibers as a collection of linear springs can reproduce bulk mechanical properties [Reinhardt and Gooch, 2018], and linear bead-spring chain models in overdamped environments show emergent viscoelasticity consistent with biological polymers [Panja, 2010]. Further, bulk mechanical properties such as strain-stiffening and compression-softening emerge from buckling and bending of individual fibers [Notbohm et al., 2015]. Taking these observations into consideration, we chose to abstract each fiber as a bead-spring chain in continuous space with linear spring potentials between direct bead neighbors and angular spring potentials between consecutive bead triplets (Fig. 1 B). This choice of mechanical model strikes a balance between simplicity and flexibility to represent buckling and bending. *In vivo* and *in vitro*, ECM fibers assemble into networks by branching and crosslinking, both of which depend on local fibril concentration as well as admixture of additional components [Kreger et al., 2010, Leonidakis et al., 2017]. To model this process, we introduced crosslinkers with linear spring potentials between beads of different fibers. Crosslinkers were added stochastically during network initialization with probability proportional to local fiber density (Supplementary Fig. 1). Movement of ECM beads was modelled by an overdamped equation of motion with added Brownian noise (Equation 7, Methods section). We implemented this model in 2D and solved the mechanical equilibrium with the high-performance molecular mechanics framework HOOMD-blue [Anderson et al., 2008, 2020].

Mechanical interaction between cells and ECM in our model occurred exclusively at spatially discrete adhesions, an abstraction for integrin adhesion complexes such as focal adhesions [Van Helvert et al., 2018]. Each adhesion in the model consisted of at least one ECM fiber bead that was mechanically coupled with a marked “adhesion site” of the CPM; adhesion beads were clamped with respect to the network dynamics, but could be moved by the cell (Fig. 1 C). During CPM simulation, different adhesion beads could become associated to the same CPM adhesion site as an abstraction for the formation of high-density integrin adhesion clusters [Michael and Parsons, 2020, Van Helvert et al., 2018]. Since only a limited amount of protein can be reasonably packed into a given volume, we set a threshold value *α*_thresh_ above which clustering additional adhesions in the same site was discouraged by an energy penalty to the cell. This model of adhesions is a natural and intuitive way to mechanically interface cell and ECM models that closely mimics integrin adhesion complexes.

The simulation was run by iteratively stepping through one Monte Carlo step (MCS) of the CPM and then one molecular dynamics (MD) integration cycle consisting of *t*_MD_ internal calculation steps of the ECM model (Fig. 1 D). We assumed that ECM network relaxation occurred on a faster timescale and was in quasi-steady state with respect to the cell. Therefore, the number of integration steps *t*_MD_ was chosen to be sufficiently large to ensure the ECM model reached mechanical equilibrium. We refer to one iteration consisting of one MCS and one MD cycle as one timestep.

### Case study: Isolated contractile cell

The observation that an isolated contractile cell deforms its surrounding ECM was a milestone experiment that launched the field of cellular biomechanics [Harris et al., 1981]. With minor alterations, this setup continues to be of fundamental importance [Malandrino et al., 2019, Mann et al., 2019, Notbohm et al., 2015, Van Helvert and Friedl, 2016]. Therefore, we simulated a purely contractile cell straining a randomly oriented fiber network as a case study to demonstrate the capabilities of the model. We envisioned the model as a thin 1 μm slice of a cell embedded in a 3D ECM network. Thus we defined a square simulation box of side length 225 μm and placed a circular cell with radius *r*_cell_ = 12.5 μm in its center. Adhesions were allowed to form in a thin 1.25 μm-wide boundary layer of the cell.

To simulate a strongly, isotropically contractile cell, we chose the CPM parameters in such a way that the cell would attempt to contract to a single lattice site. The opposing force from the ECM fiber network would counteract cell contraction. In the following, we investigate how cell-intrinsic and ECM-intrinsic parameters affect the extent of cell-induced ECM network reorganization and the extent of cell contraction, and compare these data to experiments. We focused on parameters that affect ECM network material properties: The concentration of fibers [*N*_fibers_] and of crosslinkers [*N*_crosslinkers_]; we chose values that reflected a range of sparse to dense networks. Further, we also varied the parameters affecting cell adhesion to the network: the cell’s initial adhesion density [*N*_adh_]_0_, and the threshold value for adhesion clustering *α*_thresh_. We ran simulations for 10000 timesteps (≈ 8 hours based on comparing temporal dynamics to Malandrino et al. [2019]). Unless otherwise noted, the parameter values were as listed in Table 1.

**Table 1:**
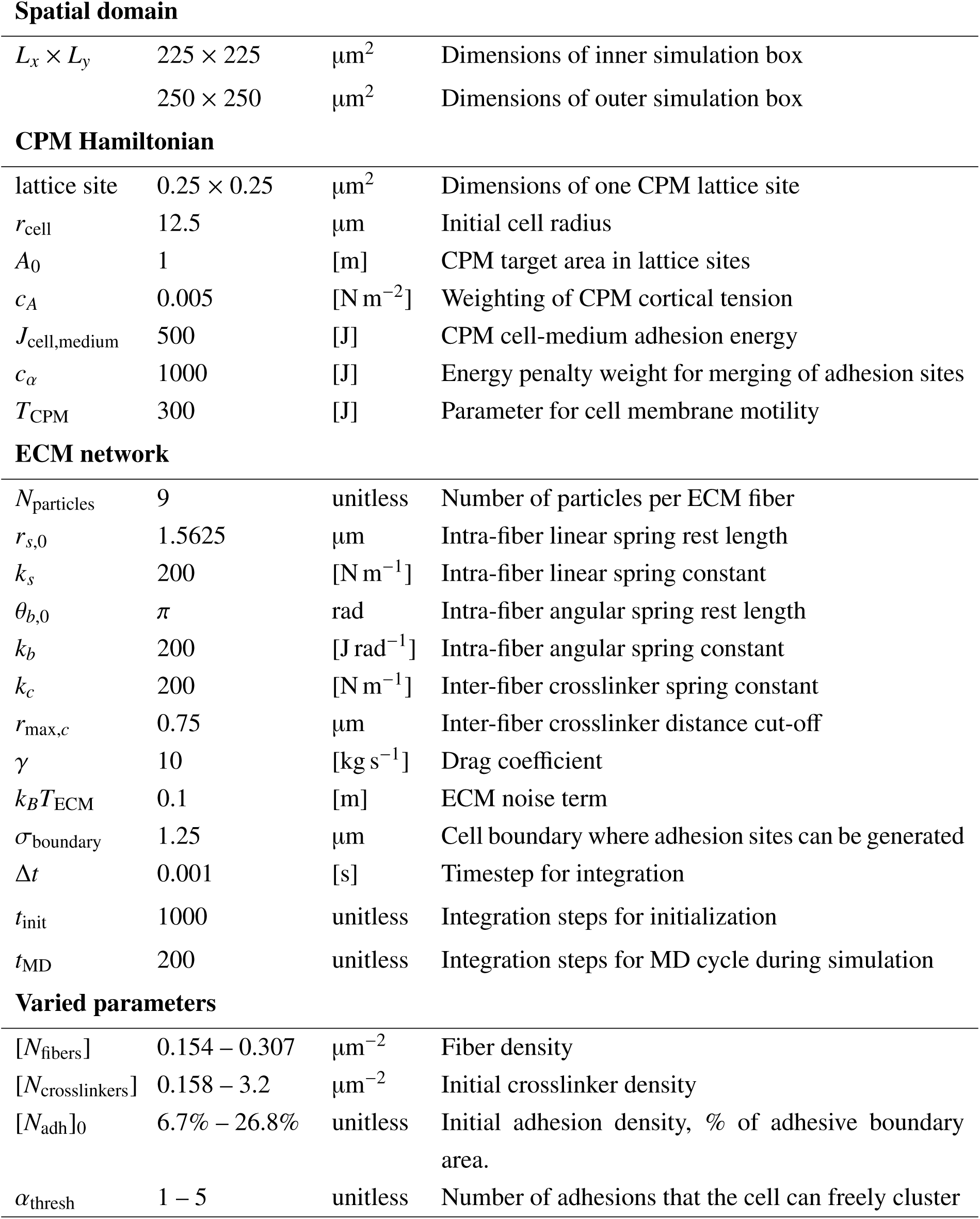
Parameter values. Square brackets show base SI units for non-dimensionalized parameters.

|  |  |  |  |
| --- | --- | --- | --- |
| **Spatial domain** |  |  |  |
| $L_x \times L_y$ | $225 \times 225$ | $\mu\text{m}^2$ | Dimensions of inner simulation box |
| | $250 \times 250$ | $\mu\text{m}^2$ | Dimensions of outer simulation box |
| **CPM Hamiltonian** |  |  |  |
| lattice site | $0.25 \times 0.25$ | $\mu\text{m}^2$ | Dimensions of one CPM lattice site |
| $r_{\text{cell}}$ | 12.5 | $\mu\text{m}$ | Initial cell radius |
| $A_0$ | 1 | [m] | CPM target area in lattice sites |
| $c_A$ | 0.005 | [N m<sup>-2</sup>] | Weighting of CPM cortical tension |
| $J_{\text{cell,medium}}$ | 500 | [J] | CPM cell-medium adhesion energy |
| $c_\alpha$ | 1000 | [J] | Energy penalty weight for merging of adhesion sites |
| $T_{\text{CPM}}$ | 300 | [J] | Parameter for cell membrane motility |
| **ECM network** |  |  |  |
| $N_{\text{particles}}$ | 9 | unitless | Number of particles per ECM fiber |
| $r_{s,0}$ | 1.5625 | $\mu\text{m}$ | Intra-fiber linear spring rest length |
| $k_s$ | 200 | [N m<sup>-1</sup>] | Intra-fiber linear spring constant |
| $\theta_{b,0}$ | $\pi$ | rad | Intra-fiber angular spring rest length |
| $k_b$ | 200 | [J rad<sup>-1</sup>] | Intra-fiber angular spring constant |
| $k_c$ | 200 | [N m<sup>-1</sup>] | Inter-fiber crosslinker spring constant |
| $r_{\text{max},c}$ | 0.75 | $\mu\text{m}$ | Inter-fiber crosslinker distance cut-off |
| $\gamma$ | 10 | [kg s<sup>-1</sup>] | Drag coefficient |
| $k_B T_{\text{ECM}}$ | 0.1 | [m] | ECM noise term |
| $\sigma_{\text{boundary}}$ | 1.25 | $\mu\text{m}$ | Cell boundary where adhesion sites can be generated |
| $\Delta t$ | 0.001 | [s] | Timestep for integration |
| $t_{\text{init}}$ | 1000 | unitless | Integration steps for initialization |
| $t_{\text{MD}}$ | 200 | unitless | Integration steps for MD cycle during simulation |
| **Varied parameters** |  |  |  |
| $[N_{\text{fibers}}]$ | 0.154 – 0.307 | $\mu\text{m}^{-2}$ | Fiber density |
| $[N_{\text{crosslinkers}}]$ | 0.158 – 3.2 | $\mu\text{m}^{-2}$ | Initial crosslinker density |
| $[N_{\text{adh}}]_0$ | 6.7% – 26.8% | unitless | Initial adhesion density, % of adhesive boundary area. |
| $\alpha_{\text{thresh}}$ | 1 – 5 | unitless | Number of adhesions that the cell can freely cluster |

### Crosslinker concentration tunes ECM network stiffness

As expected, simulated cells rearranged the ECM fiber network as they contracted (Fig. 2 A, top; Supplementary Movie 1). The ECM rearrangement was spatially heterogeneous, with areas of local fiber bunching flanked by areas of fiber thinning as highlighted by local fiber density maps (Fig. 2 A, bottom). This heterogeneity was also reflected in fiber displacement (Fig. 2 B). The cell’s ability to bundle and thus locally increase the fiber density in its surroundings depended on the crosslinker concentration (Fig. 2 C). Fiber networks with few crosslinks ([*N*_crosslinks_] = 0.32 μm^−2^) behaved as soft gels, where cells deformed the network more strongly and accumulated a “halo” of highly dense fibers (Supplementary Movie 2). Vice-versa, highly crosslinked networks ([*N*_crosslinks_] = 3.2 μm^−2^) behaved as stiff gels, having fewer dense fiber bundles and reducing overall cell contraction (Supplementary Movie 3). The qualitative difference between simulations paralleled *in vitro* experimental observations [Harris et al., 1981, Malandrino et al., 2019]. In particular, HUVEC cells embedded into fibrin gels formed a halo of dense fibers in their immediate surroundings only when the gel was polymerized with low crosslinking (Fig. 2 D; adapted from Malandrino et al. [2019]). Thus, in our model ECM stiffness emerged from network connectivity in a manner consistent with experimental data. Further, the model captured the impact of ECM stiffness on the cell’s ability to contract.

**Figure 2:**
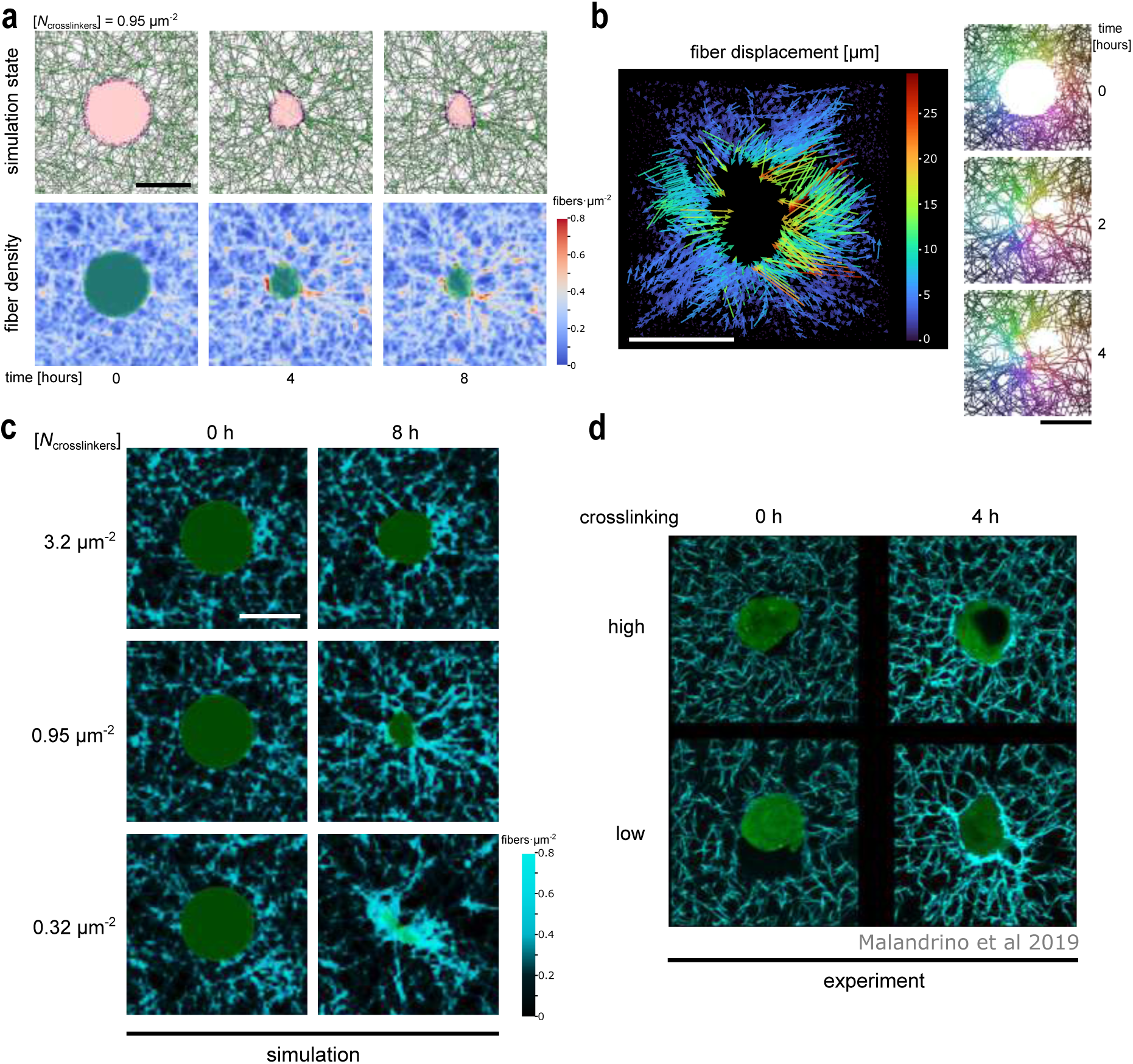
Cell contraction and ECM remodelling strongest in sparsely crosslinked networks. **a**. As the cell contracts it remodels the fiber network, creating regions of high local fiber density flanked by areas of fiber thinning. Top: Graphical output of the simulation showing CPM cell in pink, adhesion sites in purple, fibers in gray, and crosslinks in green. Bottom: Local fiber density. The cell’s position is overlayed in dark green. **b**. Left: Quiverplot illustrating fiber particle displacement due to cell contraction. Right: Three snapshots of the data displayed on the left. Each fiber is shown in a unique color. **c**. Fiber density map of initial (left) and final state (right) of simulations with different parameter values for the initial crosslinker density [*N*_crosslinkers_]. Note that the simulation for [*N*_crosslinkers_] = 0.95 μm^−2^ is the same as in panel a. We use a different color scale for better comparison to experimental data in panel d. **d**. Densification of fibers observed experimentally in HUVEC cells embedded in fibrin gels. Reproduced from Malandrino et al. [2019]. Simulation parameters in panels a and c: *α*_thresh_ = 3, [*N*_fibers_] = 0.23 μm^−2^, [*N*_adh_]_0_ = 13.4%. Scale bars: 20 μm.

### Both cell and ECM parameters affect ECM remodelling

To more precisely quantify ECM remodelling, we measured the unitless ECM densification factor *f*_dens_ and compared it with experiments. ECM densification is used to assay contractile ability, and is calculated by dividing the average fiber density near the cell by the average fiber density far from the cell (Fig. 3 A, scheme on the right). In the simulations, *f*_dens_ tended to increase over time, and the total increase depended strongly on crosslinker concentration: The densification factor increased most in soft networks with few crosslinkers, while it barely deviated from the baseline in stiff networks with many crosslinkers (Fig. 3 A). This observation in the model was consistent with *in vitro* experiments (Fig. 3 B, adapted from Malandrino et al. [2019]). Interestingly, in simulations with very low crosslinker concentration, *f*_dens_ could decrease below 1. In other words, the vicinity of the cell became more sparse. This ‘fiber thinning’ effect was due to cells pulling in loosely-connected fibers, resulting in fiber-free pockets. Fiber thinning was transient, and at longer timescales fiber densification decreased monotonically with crosslinker concentration. Therefore, for the following analyses we quantified *f*_dens_ at the final simulation state.

**Figure 3:**
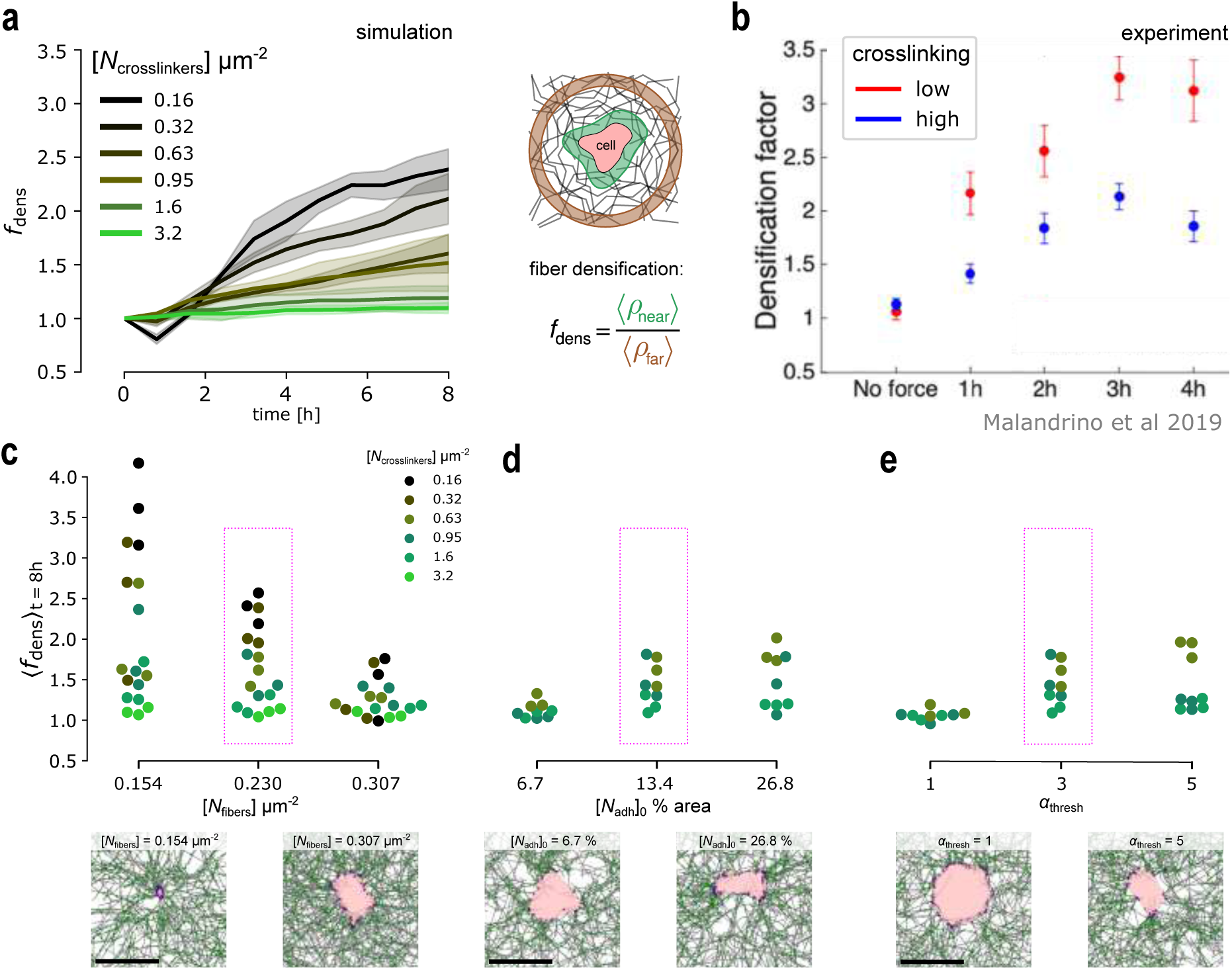
Cell adhesions, fiber density and crosslinking affect ECM remodelling. **a**. Left: Mean and standard deviation of at least 3 replicate simulations showing fiber densification over time. Simulation parameters: *α*_thresh_ = 3, [*N*_fibers_] = 0.23 μm^−2^, [*N*_adh_]_0_ = 13.4%. Right: Scheme of calculation. **b**. Experimental data showing mean and standard error of fiber densification for different levels of crosslinking. Reproduced from Malandrino et al. [2019]. **c**. Top: Effect of fiber density on average fiber densification at the final simulation state. [*N*_adh_]_0_ = 13.4%, *α*_thresh_ = 3. Bottom: Representative simulations for [*N*_crosslinkers_] = 0.95 μm^−2^. **d**. Top: Effect of initial cell adhesion density on average maximal fiber densification at the final simulation state. [*N*_fibers_] = 0.23 μm^−2^, *α*_thresh_ = 3. Bottom: Representative simulations for [*N*_crosslinkers_] = 0.95 μm^−2^. **e**. Top: Effect of maximum penalty-free adhesion clustering on average maximal fiber densification at the final simulation state. [*N*_fibers_] = 0.23 μm^−2^, *α*_thresh_ = 3. Bottom: Representative simulations for [*N*_crosslinkers_] = 0.95 μm^−2^. Note that the highlighted dataset in c, d, and e is the same, and corresponds to the data shown in a. Values for [*N*_crosslinkers_] = 0.16, = 0.32, and = 3.2 μm^−2^ were omitted in d and e. Scale bars: 20 μm.

Besides crosslinking, fiber concentration is an important modulator of network stiffness [Han et al., 2018, Steinwachs et al., 2016]. To quantify the effect of fiber concentration in our model, we compared *f*_dens_ in simulations where we varied the parameter [*N*_fibers_] (Fig. 3 C top). Networks with low fiber concentration showed a broad range of *f*_dens_ values, and were sensitive to the concentration of crosslinkers (Fig. 3 C top, Supplementary Movie 4). At high fiber concentration, the effect of crosslinker concentration diminished, likely as the networks were already sufficiently stiff to prevent cell contraction (Fig. 3 C top, Supplementary Movie 5). Consistently, an increase in either fiber or crosslinker concentration led to an increase in the effective network stiffness as quantified in *f*_dens_. In turn, effective network stiffness was inversely related to total cell contraction: In soft networks, cells were able to contract to a very small patch, while they retained a substantial area in stiff networks (Fig. 3 C bottom; Supplementary Movies 4–5).

Next, we examined how cell-intrinsic parameters affected contraction and remodelling. Cell types differ in the magnitude of traction force they exert on their surroundings [Alisafaei et al., 2021, Feld et al., 2020]. The magnitude of cellular traction force depends on various factors, including the size and stability of integrin adhesion complexes, as well as the organization and focalization of actomyosin and integrin machinery [Feld et al., 2020, Van Helvert et al., 2018]. The parameter [*N*_adh_]_0_ modelled the cell area dedicated to cell-ECM anchoring points that was initialized at simulation start. Thus, [*N*_adh_]_0_ could be interpreted as a proxy for adhesion complex size. Since all cell-ECM interactions occurred through adhesions, we expected ECM remodelling and thus densification to depend on [*N*_adh_]_0_. Indeed, when cells had very few adhesions, densification was lower, while more adhesions tended to increase densification (Fig. 3 D top; Supplementary Fig. 2 A-B). Correspondingly, cells with fewer adhesions contracted less (Fig. 3 D bottom left; Supplementary Movie 6) compared to cells with more adhesions (Fig. 3 D bottom right; Supplementary Movie 7). Thus, a larger initial adhesion area correlated with stronger effective traction force of cells in the simulation. These results correspond to the experimental observation that larger adhesions exert stronger traction forces [Van Helvert et al., 2018].

Stronger traction forces can also result from focalization of integrin clusters; cells that tend to have diffusely organized integrin contacts exert fewer forces [Feld et al., 2020, Van Helvert et al., 2018]. We modelled formation of dense integrin adhesion clusters through the parameter *α*_thresh_, which described the preference of a particular cell type to maintain several diffuse adhesions (low *α*_thresh_) or few focalized adhesion clusters (high *α*_thresh_). Reducing *α*_thresh_ resulted in cells contracting less and maintaining large areas with evenly-spread adhesions (Fig. 3 E, bottom left; Supplementary Movie 8). Conversely, increasing *α*_thresh_ resulted in cells contracting more strongly (Fig. 3 E, bottom right; Supplementary Movie 9). ECM densification also increased with increasing adhesion cluster density (Fig. 3 E top, Supplementary Fig. 2 C-D).

In summary, the cell adhesion parameters [*N*_adh_]_0_ and *α*_thresh_ modulate the effective cell traction forces that emerge from the mechanical interactions between the fiber network and the contracting cell. Furthermore, the qualitative impact of these parameters on ECM remodelling and cell contraction is consistent with experimental observations.

### Long-ranged force propagation depends non-linearly on crosslinkers

The ability of fibrous ECM to transmit long-range forces distinguishes it from synthetic substitutes [Wang and Xu, 2020]. To assay how cell contraction propagated through the ECM network in the model, we calculated the average displacement of fibers in a radius of 30 μm centered on the cell. Interestingly, the displacement at the final simulation state was non-linear with respect to crosslinker density (Fig. 4 A, Supplementary Fig. 3 A-B). The initial increase in displacement with crosslinker density was related to fiber thinning in sparsely-crosslinked networks which we also noted earlier. Intuitively, in the extreme scenario of a completely unconnected network, only fibers that are directly attached to the cell will displace, and the cell’s force is not propagated at all into the network. Indeed, at the lowest crosslinker concentration of 0.16 μm^−2^, most of the fibers are not connected and fiber-free pockets form as the cell contracts (Fig. 4 A, B left; Supplementary Movie 10). Past a certain minimum connectivity, increasing the crosslinker density simply stiffened the network, increasing resistance to cell contraction and thus reducing displacement. At [*N*_crosslinkers_] = 0.63 μm^−2^ we obtained the maximum displacement in terms of magnitude (Fig. 4 A, B middle; Supplementary Movie 11). Long-ranged force propagation still occurred at higher crosslinker concentrations, but the magnitude of fiber displacement decreased (Fig. 4 A, B right; Supplementary Movie 12). With the exception of sparsely-connected networks, displacement magnitude followed the same trends as fiber densification with respect to parameter changes, being greatest when fiber density was low and cells were strongly contractile (high initial adhesion density or high clustering threshold; Supplementary Fig. 4, 5, 6).

**Figure 4:**
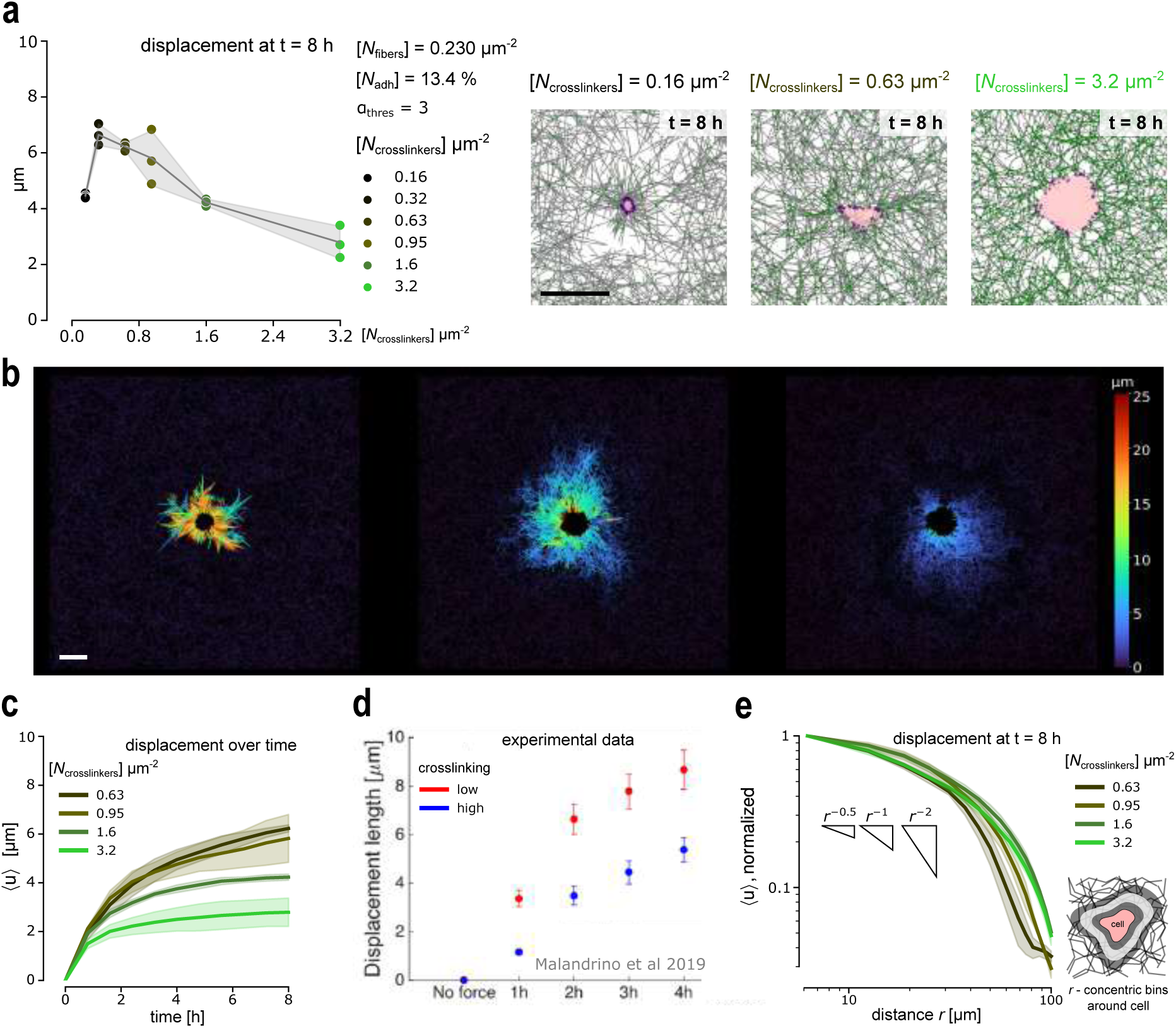
Long-ranged force propagation requires highly-crosslinked networks. **a**. Left: Absolute average fiber displacement as a function of crosslinker concentration. Plotted are individual simulation data (points), and their mean and standard deviation. Right: Simulation state snapshots showing the extent of cell contraction and fiber displacement. Note fiber thinning due to loosely-connected fibers at the lowest concentration. **b**. Quiverplots showing total displacement with respect to the initial state for the three simulations shown in a. **c**. Total fiber displacement as a function of time. Displacement magnitude was averaged in a radius of 30 μm around the center of the simulation domain; mean and standard deviation of at least three simulations each. Simulation parameters: [*N*_fibers_] = 0.23 μm^−2^, [*N*_adh_]_0_ = 13.4%, *α*_thresh_ = 3. **d**. Experimental data showing mean and standard error of ECM displacement for different levels of crosslinking. Reproduced from Malandrino et al. [2019]. **e**. Average fiber displacement at equilibrium decayed nonlinearly with distance from the cell. After normalization, curves displayed similar dynamics for [*N*_crosslinker_] ≥ 0.63 μm^−2^. Displacement was averaged in concentric areas from the cell edge at t = 8 h; mean and standard deviation of at least three simulations each. Simulation parameters: [*N*_fibers_] = 0.23 μm^−2^, [*N*_adh_]_0_ = 13.4%, *α*_thresh_ = 3. Scale bars: 20 μm.

We next focused on simulations where [*N*_crosslinkers_] ≥0.63 μm^−2^, since this parameter range enabled long-ranged force propagation. Cell contraction and cell-induced displacement of nearby ECM fibers proceeded in two phases: A first phase lasting roughly 1 hour with a large rate of displacement was followed by a second longer phase with a lower rate of displacement that converged to an equilibrium in simulations with stiffer ECM networks (Fig. 4 C, Supplementary Fig. 3 C-D). The shape of the displacement curves qualitatively matched experimental observations: contractile cells provoked large ECM strains at a timescale of 1-2 hours and displacement slowly converged to a plateau over a timescale of 4 hours (Fig. 4 D; reproduced from Malandrino et al. [2019]).

The average fibre displacement ⟨*u*⟩ decays with increasing distance *r* from a contractile cell’s edge with the power law ⟨*u*⟩ = *r*^−*n*^ [Wang and Xu, 2020]. In an ideal isotropic linear elastic material such as polyacrylamide gels, the exponent is given by *n* = 2 in 3D and *n* = 1 in 2D, that is respectively ⟨*u*⟩ = *r*^−2^ or ⟨*u*⟩ = *r*^−1^. In contrast, anisotropic materials such as gels constituted of ECM fiber networks have *n* < 1 up to a certain threshold distance, where the exponent *n* increases again [Wang and Xu, 2020]. Intuitively, the force exerted by the cell is transmitted up to a certain distance in the network by “force chains” – collections of mechanically crosslinked fibers. We quantified the displacement-decay relation in our simulations by averaging the fiber displacement in increasing concentric bins from the cell edge in the final simulation state. When these data were rescaled such that the maximum average displacement was set to 1, we observed that the resulting curves nearly overlapped (Fig. 4 E, Supplementary Fig. 3 E-F). As described theoretically [Wang and Xu, 2020], we observed a region with a decay exponent of *n* ≈ 0.5 close to the cell that transitioned to a region with *n* > 1 far from the cell. Interestingly, the decay exponent of *n* ≈ 0.5 matched experimentally measured values near cells embedded in fibrin gels [Notbohm et al., 2015]. Together, these results demonstrate that our model captures long-ranged force transmission, a typical mechanical feature of ECM fiber network gels.

### Cell-induced local strain-stiffening of the ECM

Fibrous biopolymer gels stiffen when strained, which is thought to contribute to mechanosensing and mechanotransduction in multicellular pattern formation [Van Helvert and Friedl, 2016, Wang and Xu, 2020]. Local stiffening of the ECM by cell-generated strains has been measured using nanoindentation with atomic force microscopy (AFM) [Van Helvert and Friedl, 2016]. In this application of AFM, a spherical probe was pressed with a predefined force into the ECM; the resulting force-distance relation was used to determine the ECM’s local stiffness given by Young’s modulus [Van Helvert and Friedl, 2016]. Inspired by these experiments, we implemented AFM *in silico* using the MD part of the model (Fig. 5 A; see Methods section for a detailed description). Briefly, we introduced a new particle to simulate the AFM probe and pressed it with constant force in the z-direction on the ECM network. The AFM probe particle and ECM network particles interacted with a Lennard-Jones potential. A grid of linear elastic pillars was placed underneath the 2D ECM thin film in order to mimic an extended medium in the third dimension. Thus, in regions with few fibers only the pillar grid provided a counterforce to probe indentation. In regions with many fibers, probe indentation was resisted by the combination of the pillar grid and the overlying fiber network.

**Figure 5:**
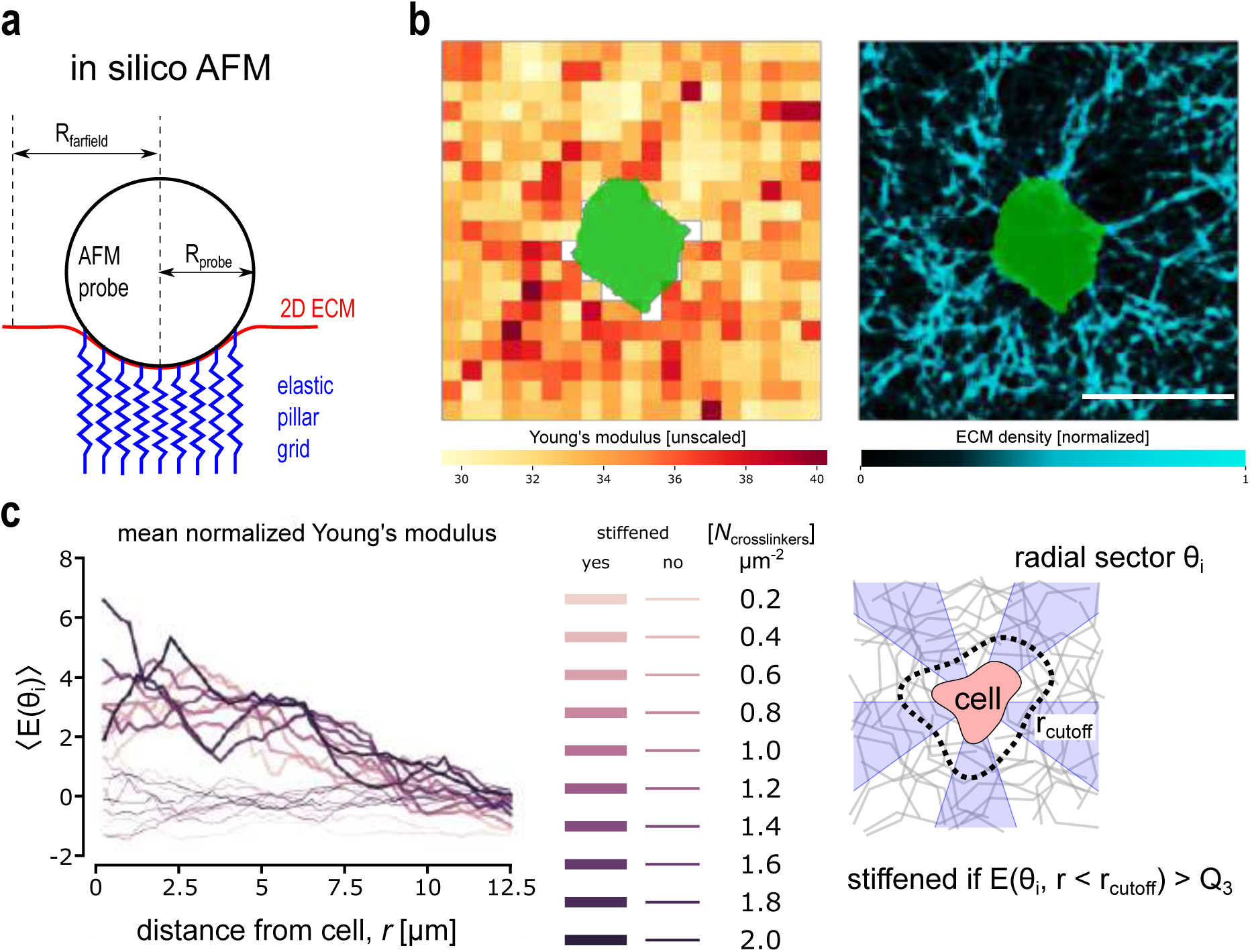
In silico AFM shows local stiffening in areas of fiber densification. **a**. Schematic representation of the in silico AFM setup. **b**. Left: Heatmap of measured Young’s moduli of a representative simulation at t=4 h. Right: Fiber density map with values normalized to the interval [0, 1]. **c**. Mean background-corrected Young’s moduli ⟨*E*⟩ as function of distance from the cell in radial sectors. Background correction was done by subtracting the mean Young’s modulus of the entire domain before cell contraction (i.e. when t=0 h). Stiffened sectors are defined as those sectors *i* where ⟨*E*(*θ*_*i*_)⟩ within 5 μm from the cell perimeter > 75% quantile of all sectors. The unstiffened sectors are plotted with thinner lines. Each pair of thick and thin lines correspond to one simulation with parameter values *a*_thresh_ = 3, [*N*_fibers_] = 0.1536 μm^−2^, [*N*_adh_]_0_ = 13.4 %. Scale bars: 20 μm.

First, we validated the *in silico* AFM methodology in cell-free networks. We measured Young’s modulus in cell-free isotropic networks of varying fiber and crosslink density at various levels of externally applied prestrain, and using various choices of pillar grid elasticity *k*_pillar_ (Supplementary Fig. 7). This parameter scan revealed that Young’s modulus depended linearly on fiber density, while the elasticity of the pillar grid only introduced a linear offset according to the following linear relation:

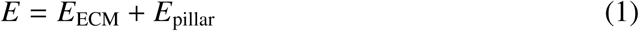

where *E* is the Young’s modulus determined via AFM, *E*_pillar_ is the modulus of the pillar grid, and *E*_ECM_ is the contribution due to the 2D ECM, which followed the power law relation

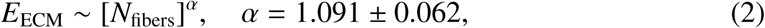

We verified the validity of this power law for strain values up to 20%. Increasing the externally applied prestrain shifted the distribution of Young’s moduli to larger values (Supplementary Fig. 7). Thus, our fiber network replicated strain-stiffening under externally-applied strains.

Next, we applied *in silico* AFM to our simulations of a contracting cell in a fiber network. Using the equilibrium configuration of the network at t = 4 h, we applied the virtual probe at regularly spaced intervals. The resulting spatial maps of Young’s modulus showed that local stiffness correlated to local fiber density (Fig. 5 B). To more precisely quantify the extent of cell-induced strain-stiffening, we measured Young’s modulus as a function of the distance *r* from the cell perimeter (Fig. 5 C). To account for the fact that cell contraction created areas of local fiber bunching flanked by areas of fiber thinning, we subdivided the cell’s surroundings into sectors of angle *θ* relative to the horizontal axis crossing the cell centroid (Fig. 5 C, scheme on the right). For each sector *θ*_*i*_, we computed the mean Young’s modulus ⟨*E* (*θ*_*i*_)⟩ as a function of *r*. We background-corrected these data by subtracting the mean background modulus of the network in the absence of cell-applied forces. Thus, ⟨*E* (*θ*_*i*_)⟩(*r*) ≈ 0 in areas without stiffening, while ⟨*E* (*θ*_*i*_)⟩(*r*) > 0 if strain-stiffening has occurred. Consistent with our prior observations that cells densified fibers in their immediate surroundings, we observed maximal stiffness within 5 μm from the cell perimeter. Thus, we chose the cutoff distance *r*_cutoff_ = 5 μm to define “stiffened” sectors as sectors where ⟨*E* (*θ*_*i*_)⟩(*r* < *r*_cutoff_) was above the 75% quantile of all measured sectors in one simulation run. Other sectors were defined as “unstiffened”. As expected, the average ⟨*E* (*θ*_*i*_)⟩ in unstiffened sectors was ≈ 0 regardless of distance to the cell (Fig. 5 C, thin lines). In contrast, the average ⟨*E* (*θ*_*i*_)⟩ in stiffened sectors was clearly > 0 close to the cell, indicating cell-induced strain-stiffening (Fig. 5 C, thick lines). Far from the cell, stiffened sectors reduced to ⟨*E* (*θ*_*i*_)⟩ ≈ 0, as expected since the cell’s pull only propagated up to a certain distance in the network. The difference between stiffened and unstiffened sectors held true regardless of crosslinker density (Fig. 5 C, shades of color), similar to what we observed for steady state fiber displacement (Fig. 4 E). In summary, our data show that cell-mediated fiber densification locally increases the network stiffness.

## Discussion

Cell-ECM mechanobiology is fundamental to morphogenesis, homeostasis, growth and regeneration [Sree and Tepole, 2020, Walma and Yamada, 2020]. An outstanding challenge in the field is relating measurable macroscale mechanics to microscale properties of cells and ECM cross-talk [Sree and Tepole, 2020]. ECM fiber network models can adequately address this challenge in sparse networks [Eichinger et al., 2021, Wang and Xu, 2020] or when populations at high cell density can be approximated with a continuum [Guo et al., 2022]. When individual cell behavior and ECM fiber dynamics are non-negligible, a more detailed model of both is needed [Eichinger et al., 2021, Guo et al., 2022, Sree and Tepole, 2020]. We here provide such a model by introducing a coarse-grained discrete mechanical model of ECM fibers hybridized with CPM cells, and demonstrate the potential to calibrate it to biomechanical measurements, e.g. with AFM. We recreated a classic experimental setup of an isolated contractile cell embedded in an ECM fiber network, and showed that various metrics – including fiber densification, long-ranged force propagation with nonlinear displacement-decay, and local cell-induced stiffening – agreed with experimental findings. These results emerged from dynamic interactions between the contracting cell and the fiber network, and were not explicitly coded into the model. Thus, our model captures essential features of cell-ECM mechanobiology. We expect that further refinement in fiber network construction heuristics based on measured network properties will improve the quantitative agreement between simulations and experimental data.

Cells in ECM fiber networks exhibit different biomechanical behavior compared to cells in synthetic gels such as polyacrylamide [Notbohm et al., 2015, Rudnicki et al., 2013]. This discrepancy has been variously ascribed to the length scale of fibers with respect to the size of the cell, the fiber structure, the network topology, or the ability of fibers to buckle or bend [Davidson et al., 2019, Notbohm et al., 2015, Ronceray et al., 2016, Wang and Xu, 2020]. In our model, we used an abstract fiber network whose structure and topology were not calibrated to experimental data. Yet our simulation produced qualitatively comparable results to experiments. Note that our bead-chain model in principle permits fiber buckling and bending, but they were not energetically favored with the parameters we used. Therefore, our model suggests that for the biomechanical properties quantified here, the length scale of fibers with respect to the cell plays the major role. Theoretical work supports this conclusion, in that fiber networks with fundamentally different topologies show qualitatively similar mechanical properties, and differ mostly quantitatively in the spatial heterogeneity of the response [Humphries et al., 2017]. Long-ranged force transmission through a network is also largely independent of the precise network structure [Humphries et al., 2017]. A feature characteristic of long-ranged forces in biological ECM is the formation of ‘mechanical bridges’ macroscopically visible as fiber bundles [Notbohm et al., 2015]. We have observed fiber bundling typical of mechanical bridges between cells (Fig. 6; Supplementary Movie 13). We will investigate this finding and its impact on multicellular pattern formation more deeply in future work.

**Figure 6:**
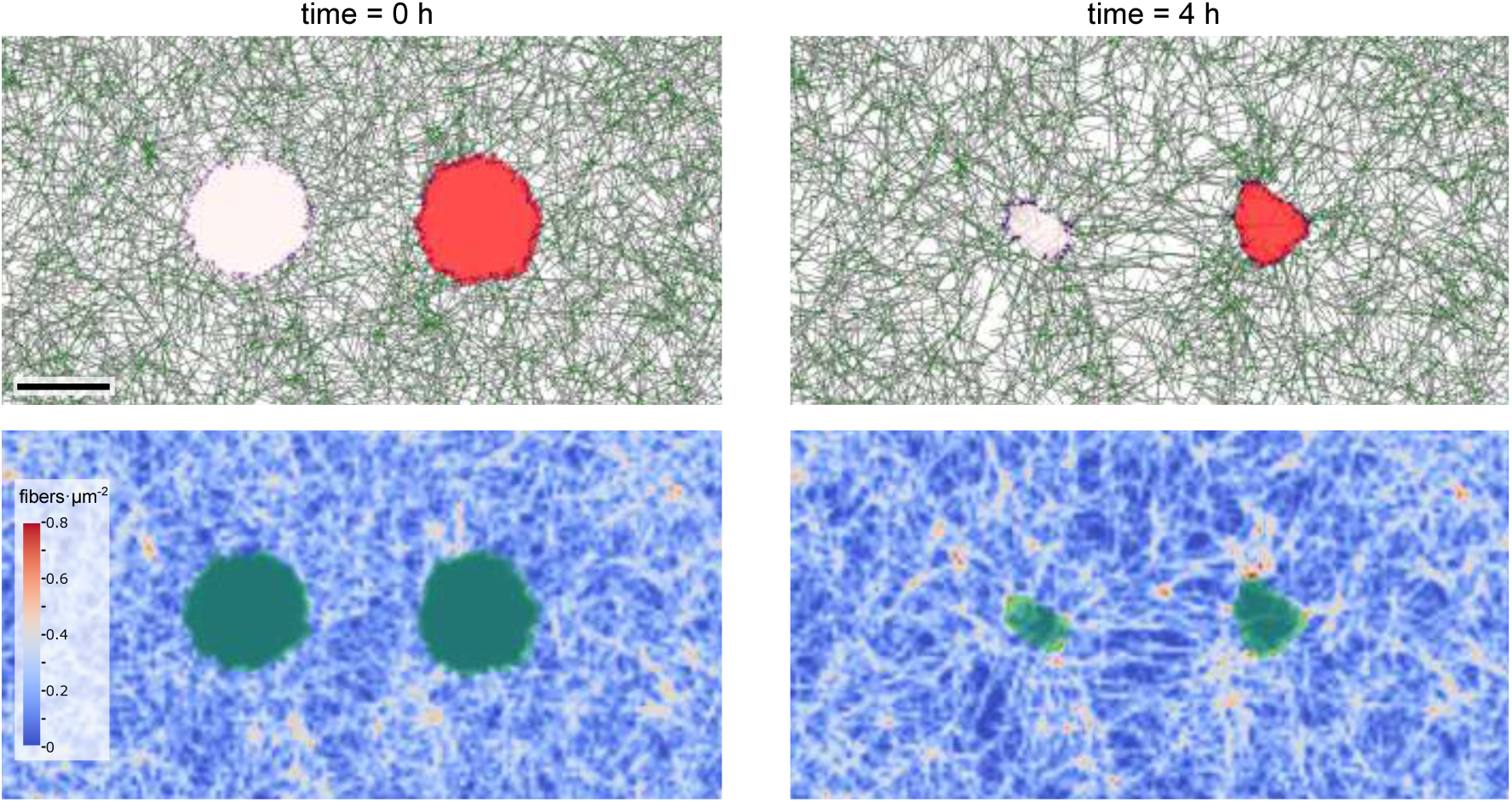
Fiber bundles form between two contractile cells. Top: Initial and final simulation state. Bottom: Fiber density map. Scale bar: 20 μm.

Cell contraction is a comparatively ‘simple’ cell behavior. Although the contractile cell could also be represented entirely using the molecular dynamics model (HOOMD-blue), e.g., by using the subcellular element approach [Newman, 2005, Sandersius and Newman, 2008], our hybrid approach will show its strength particularly in more complex problems. The hybrid approach can benefit from an enormous ‘library’ of available extensions of the CPM for modelling complex cell behavior ranging from chemotaxis [Savill and Hogeweg, 1997] and anomalous cell migration [Niculescu et al., 2015, Van Steijn et al., 2022], to complex multicellular developmental mechanisms such as angiogenesis [Van Oers et al., 2014] and somitogenesis [Hester et al., 2011, Nelemans et al., 2020]. Network structure likely plays a prominent role in such complex cell behavior. For example, during collective cell migration, so-called ‘leader’ cells reorient ECM fibers as they migrate, creating a path of least resistance for ‘follower’ cells to migrate through [Van Helvert et al., 2018]. Extending our model to include complex (multi-)cell behavior will be the focus of future work. This would entail incorporating dynamic adhesion turnover, e.g. as done in a previous CPM featuring a continuum description of ECM mechanics [Rens and Merks, 2020], as well as enabling cells to push on fibers during migration by volume exclusion, a hallmark of cell migration in ECM-dense environments [Van Helvert et al., 2018]. Such model extensions will also require the simulation to be scaled up; particularly the increase in number of fibers might pose computational challenges, which have hampered previous approaches to modelling cell-ECM mechanobiology [Reinhardt and Gooch, 2014, 2018]. For this reason, we specifically chose to work with the established and optimized library HOOMD-blue, which supports GPU-based computing [Anderson et al., 2008, 2020]. We expect that performance can be improved even further by reducing the ratio between MCS and MD steps comprising one timestep. These optimizations will enable to scale simulations to analyze the role of cell-ECM mechanobiology in the collective behavior at the mesoscale of dozens of cells.

## Methods

### Computational model

We interfaced a bead-chain model for ECM fiber networks with a CPM representing the cell. A high-level overview of initialization and simulation procedures is given in Fig. 7. In the following sections we explain in detail how the fiber network was initialized and integrated, how the CPM was initialized and interfaced to the fiber network, which terms were included into the CPM Hamiltonian, and how the simulation was performed.

**Figure 7:**
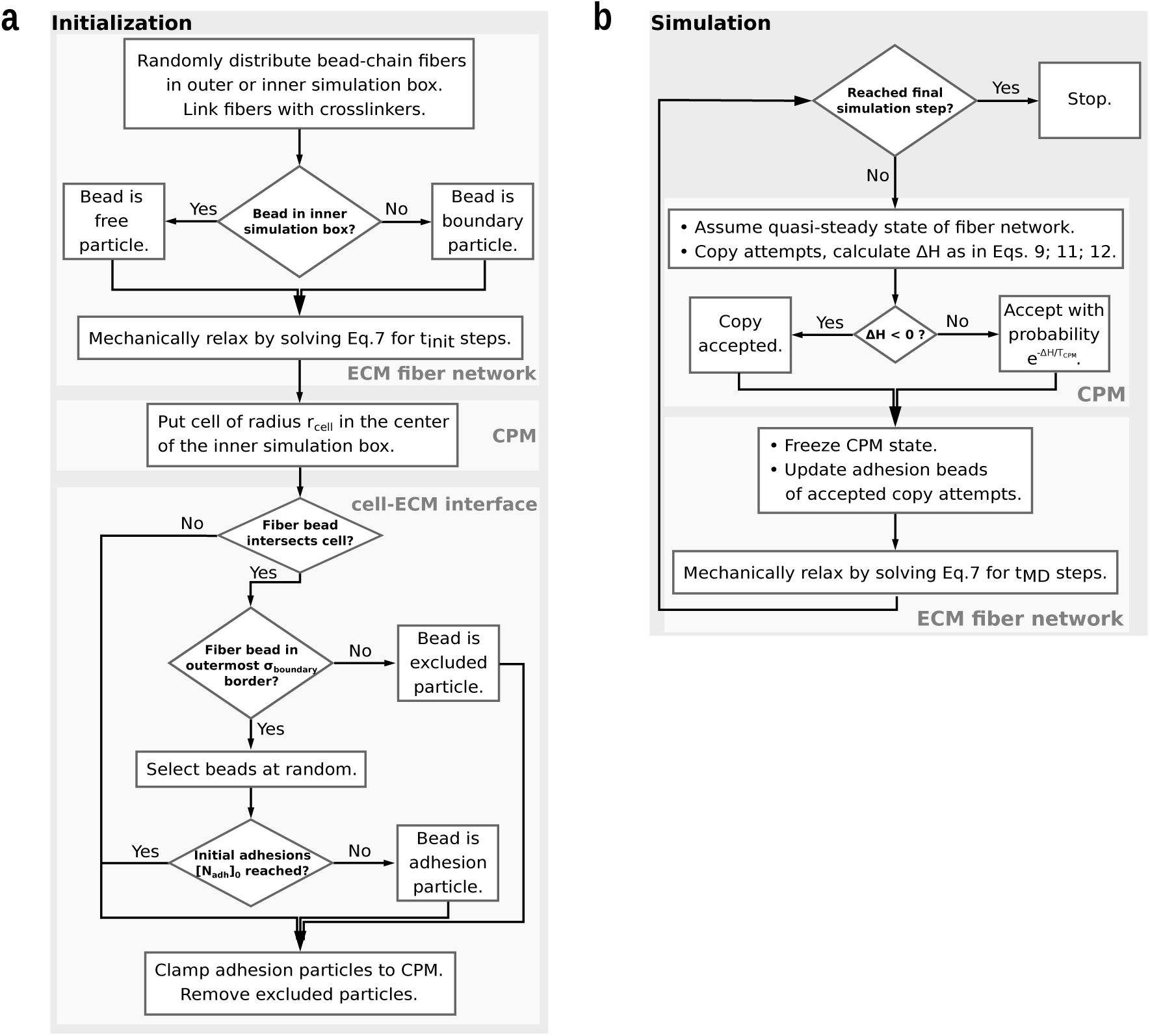
Flowchart illustrating the model. **a**. Initialization procedure. **b**. Simulation procedure.

### ECM fiber network generation

The spatial domain was set to a 2-dimensional rectangle with dimensions (*L*_*x*_ + 2*h*) × (*L*_*y*_ + 2*h*) (“outer simulation box”), where *h* = (*N*_particles_ − 1)*r*_*s*,0_ with *r*_*s*,0_ was the total rest length of each fiber with *r*_*s*,0_ the rest length of the linear spring used to model intra-fiber particle interactions. Centered in this domain we defined a rectangular “inner simulation box” with dimensions *L*_*x*_ × *L*_*y*_. The outer simulation box was chosen in such a way to ensure that the average fiber concentration was constant in the inner simulation box. We seeded a number of *N*_fibers_ of bead-chain fibers with the same number *N*_particles_ in the outer simulation box. For each fiber, we first placed an initial bead at a random position **x**_0_ uniformly within the outer simulation box. To place subsequent beads in the chain, we chose a random unit vector 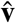 uniformly once for each fiber. Then, a new bead position was found by computing 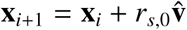.

After construction of the fibers, crosslinkers were added to form a network. The crosslinkers were stochastically distributed according to the ‘local fiber density’. We computed the local fiber density by binning the inner simulation box into equally-sized square bins, and counting for each bin *k* the number of unique fibers *f*_*k*_ passing through it and its Moore neighborhood (Supplementary Fig. 1). We then sampled *N*_crosslinks_ bins with probability proportional to the local fiber density: 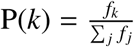. In each sampled bin, we selected a pair of fibers (*i, i*′), *i* ≠ *i*′ uniformly at random from the unique fiber pairings possible within the Moore neighborhood of the sampled bin. We computed the probability of adding a crosslinker between all possible bead pairs (*x*_*i,n*_, *x*_*i*′_,*m*); *n, m* ∈ [1, *N*_particles_] of the sampled fibers by

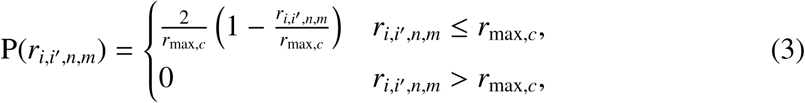

where *r*_*i,i*_, *n,m* = ‖*x*_*i,n*_ − *x*_*i*′, *m*_‖ was the distance between the two beads in a pair. A crosslinker was then added between one of the pairs of beads (*x*_*i,n*_, *x*_*i*′, *m*_) with probability 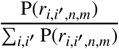. The crosslinker spring’s rest length was set to be equal to *r*_*i,i*′, *n,m*_. Note that if *r*_*i,i*′, *n,m*_ > *r*_max,*c*_ for all pairs of beads, then no crosslink was added.

### Mechanical model and integration of the ECM fiber network

Each fiber was made up of *N*_particles_ beads 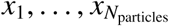. Consecutive beads in fibers were connected via Hookean springs, resulting in the potential energy

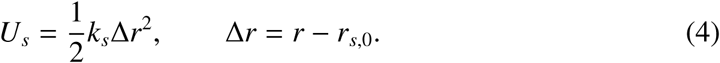

Here *k*_*s*_ was the spring constant, and Δ*r* was the deviation from the resting length *r*_*s*,0_. Similarly, crosslinkers were modelled as Hookean springs whose rest length *r*_*c*,0_ was set to be equal to the initial distance between crosslinked beads determined during network generation (see previous section)

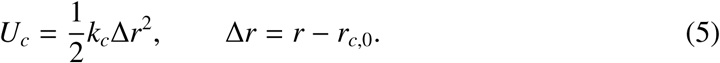

Furthermore, we included angular springs between three consecutive beads in a fiber, with potential energy

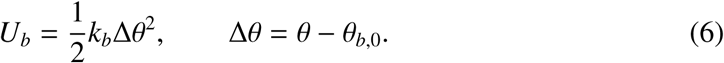

Here *k*_*b*_ was the bending rigidity, and Δ*θ* was the deviation from the resting angle *θ*_*b*,0_, and *θ* was the interior angle between consecutive bead triplets along the fiber.

Inertial effects were neglected, and thus the dynamics were governed by the over-damped Langevin equation

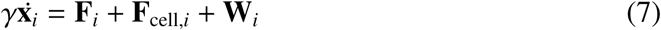

where γ was the drag coefficient, **F**_*i*_ was the mechanical force from the fiber network, which included extension and bending of the fibers and extension of the crosslinks, **F**_cell,*i*_ was the loading force generated by the cell (explained in the next section), and **W**_*i*_ was a random force satisfying

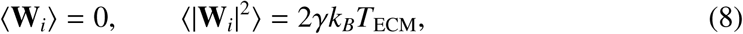

with *k*_*B*_ the Boltzmann constant and *T*_ECM_ a parameter that set the degree of noise in the system. Dirichlet boundary conditions were enforced by clamping beads outside the inner simulation box. Integration of this system was done using the HOOMD-blue molecular dynamics library [Anderson et al., 2008, 2020].

### Extended cellular Potts model

We adapted the cellular Potts model (CPM) from previous work [Daub and Merks, 2013, 2014]. The CPM lattice was contained entirely in the inner simulation box, which we subdivided with a regular square lattice with side length 0.25 μm; the outermost row of lattice sites was made impassable. Cells were represented by a collection of lattice sites *p* with equal indices *σ*_*p*_. The change in spatial configuration of lattice sites required physical work, described by a Hamiltonian energy function *H*. The term describing the energy of a particular spatial configuration was

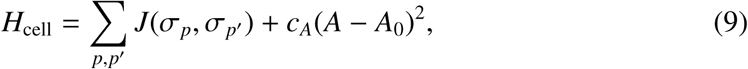

where the sum was over neighboring lattice sites (*p, p*′) with *σ*_*p*_ ≠ *σ*_*p*′_, *J* modelled adhesion energies of interfaces between different cell types or the medium, *c*_*A*_ corresponded to cell contractility, *A* was the cell area during simulation runtime, and *A*_0_ the target area.

The cell was coupled to the ECM via ‘adhesions’, which represented cross-membrane complexes through which the cytoskeleton can form mechanical bonds with the ECM. Adhesions were modelled as beads of the ECM which were associated to a marked lattice site in the cell. During an MD cycle, these adhesion-associated beads were mechanically clamped, and were thus exempted from integration of Equation 7. They were specified at initialization of the simulation by sampling *N*_adh0_ ECM beads which intersected with a narrow outer boundary of the initial cell configuration; the width of this boundary was set to *σ*_boundary_ = 1.25 μm.

We assumed that the ECM was in quasi steady state. As such, the net force on each of the adhesions was 0 N, hence for the force on an adhesion-associated bead *x*_*i*_ we obtained the balance equation

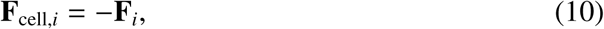

where **F**_*i*_ was the opposing mechanical force from the fiber network. Suppose lattice site *p* contained an adhesion-associated bead *x*_*i*_. A CPM copy attempt of a neighboring lattice site *p*′ onto site *p* would result in the displacement of this bead onto a new lattice site 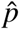, which was chosen to be a random neighbor of *p* belonging to the same cell *σ*_*p*_. The work the cell had to exert on the ECM for this displacement was then, in light of the balance equation,

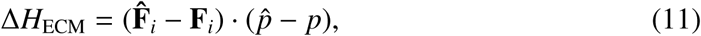

where 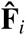 denoted the mechanical force from the ECM after the displacement of bead *x*_*i*_ in direction 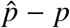. This force was obtained from Equations 4, 5 and 6 taking only bead *x*_*i*_’s direct neighbor beads into account. A copy attempt thus involved the selection of a neighbor pair (*p, p*′) with *σ*_*p*_ ≠ *σ*_*p′*_, as well as displacement of adhesion beads associated with *p* to 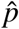 with 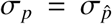. If no valid neighbor site 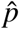 existed, then the adhesion site was annihilated and all beads associated with *p* were released from mechanical clamping, turning to ‘free’ particles subject to Equation 7. If both sites 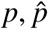 contained adhesion-associated beads, they were allowed to cluster, simulating the creation of a larger adhesion complex from smaller ones. Since there is a physical limitation on the amount of adhesion proteins which can fit into a given part of the cell, we introduced a penalizing energy term associated with the local density of adhesion-associated beads of the form

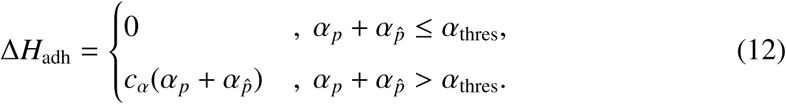

Here, sites 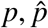 were neighbors belonging to the same cell; *α*_*p*_ and 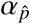 were, respectively, the adhesion density in lattice sites 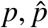; and *c*_*α*_ and *α*_thresh_ were model parameters. The parameter *c*_*α*_ was a weighting factor for the penalty term, while *α*_thresh_ was the threshold of adhesion density above which the penalty was applied. Hence a copy attempt which would result in fusion of adhesions from sites *p* and 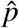 past the threshold value *α*_thresh_ received an energy penalty.

The total work required for a copy attempt was then the sum of the terms in Equations 9, 11, and 12:

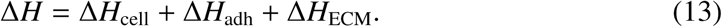

This free energy governed the dynamics of the cell by means of a Metropolis-type algorithm. The copy attempt was always accepted if Δ*H* < 0 and accepted with probability 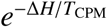 if Δ*H* ≥ 0, where the reference energy *T*_CPM_ represented cell motility.

### Simulation algorithm

To model the dynamic interaction between the cell remodelling the ECM by means of contractive force, and the effect of ECM structure on the cell dynamics, we first initialized the network and CPM cell, then iteratively applied the two models we just described (see Algorithm 1; Fig. 7).

#### Algorithm 1: Simulation algorithm

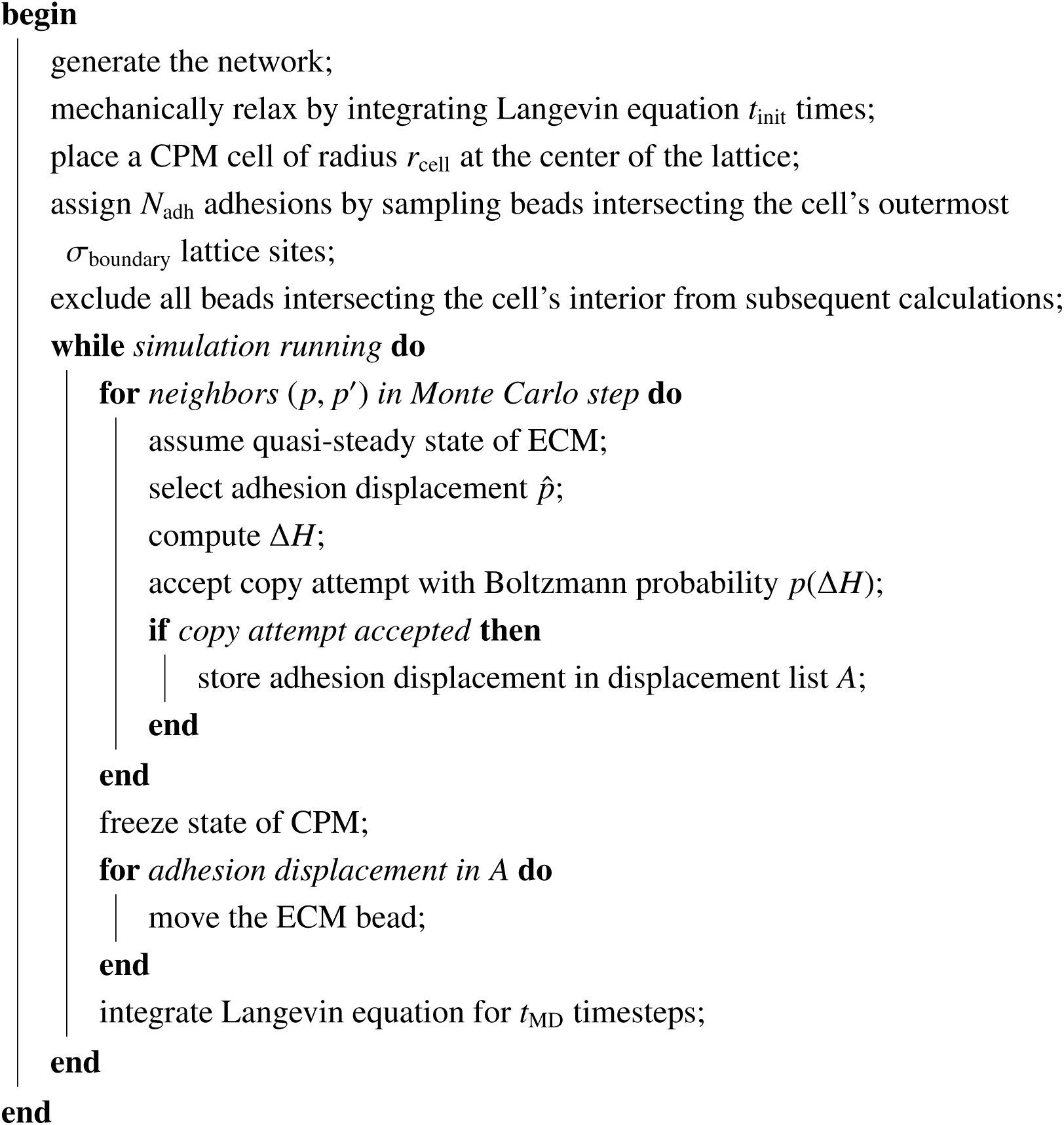

#### Parameter values

Unless otherwise noted, parameter values were as listed in Table 1. We estimated the spatial and temporal scale of the model based on data in Malandrino et al. [2019]. Specifically, we used the size of Cytochalasin D-treated cells to assign an uncontracted cell radius of *r*_cell_ = 12.5 μm. For the temporal scale, we used the dynamics of ECM fiber densification and displacement after release of Cytochalasin D inhibition to arrive at an estimate of 5000 simulation timesteps ≈4 h; for most parameter combinations the model reached equilibrium within this time frame. Force and energy-related parameters are non-dimensionalized.

We varied the ECM fiber density in the range of 0.154 – 0.307 μm^−2^; crosslinkers, which represented additional linkage fibrils, were varied in the range of 0.158 – 3.2 μm^−2^. These values resulted in a range of sparse and unconnected networks to very dense highly-connected networks. Considering we modelled a 1 μm tall thin slice of a 3D volume, and assuming that fibers were cylindrical with 0.25 μm diameter and total rest length of (*N*_particles_ − 1)*r*_*s*,0_ = 12.5 μm, we obtained a range of 9.42 – 18.85 % volume fraction of fibers. Analogously for crosslinkers, if we considered crosslinker rest length as the maximum *r*_*max,c*_, we obtained a range of 0.58 – 11.78 % volume fraction of additional fibrils. Note that this calculation is an upper bound as not all crosslinkers may successfully form, and they may form at shorter rest lengths. Therefore the total range of fiber plus crosslinker density assayed in simulations was 10 – 30.63 % volume fraction. For comparison, collagen gels *in vitro* have typical ranges of 7.5 – 18 % volume fraction (or, equivalently 0.5 – 3 mg mL^−1^) [Kreger et al., 2010].

The concentration of initial adhesions [*N*_adh_]_0_ was varied in the range of 6.7% – 26.8% area fraction of the adhesive cell boundary, which for the parameters we used corresponded to an absolute number of 100 – 400 adhesions. Note that, as with crosslinkers, these are upper bounds, since adhesions could only form if fibers were within the adhesive boundary of the cell in the initial condition. Thus, in sparse networks fewer adhesions may form. During simulation runtime, adhesions were allowed to cluster up to *α*_thresh_ times, where *α*_thresh_ was varied in the range 1 – 5. Unless otherwise noted we used [*N*_adh_]_0_ = 13.4%, *α*_thresh_ = 3, resulting in roughly 66 adhesion clusters being energetically favorable. For comparison, mature focal adhesions in MDA-MB-231 fibroblasts and in endothelial colony-forming cells were estimated to be in the range of 20 – 60 [Horzum et al., 2014, Mason et al., 2019], and in some experimental conditions up to 125 [Mason et al., 2019].

### In silico AFM

In order to quantify cell induced strain stiffening of the ECM, we implemented *in silico* atomic force microscopy (AFM) using molecular dynamics. Since our model represented the ECM as a 2D thin film, we introduced an artificial isotropic third dimensional component by placing a grid of linear elastic pillars underneath the ECM (see also Fig. 5 B). A spherical probe of radius *R*_probe_ was then positioned on top of the ECM; areas of 6.25 μm^2^ were probed at a minimum distance of 1 μm from the cell. The probe interacted with both the ECM thin film and the pillar grid via a truncated Lennard-Jones potential. Varying forces *F*_probe_ were applied at the probe using molecular dynamics, and the corresponding probe deflection δ_probe_ was recorded. Since the induced deformation of ECM remained localized, the portion of the ECM which was integrated was restricted to particles lying within a distance *R*_farfield_ from the probing site.

To relate the obtained force-deflection curve to the Young’s modulus, we used Hertz’ law

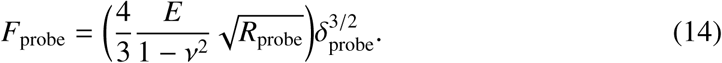

Here ν was the Poisson ratio and *E* was the Young’s modulus. Note that: (1) this model assumed infinite sample thickness, which we approximated using the pillar grid; (2) this model assumed linear isotropic elasticity, not taking into account the complex stress-strain behaviour which may emerge from collagen networks; (3) although the Poisson ratio of soft biological tissue was often assumed to be 0.5 [Lin et al., 2007], for collagen gels it may be different and spatially varying [Steinwachs et al., 2016]. To prevent these limitations from affecting our results, we obtained *E* from (14) using a linear regression fit, and discarded measurements for which *R*^2^ < 0.995, similar to Van Helvert and Friedl [2016].

**Table 2:** Parameter choices for the AFM model. Square brackets show base SI units for non-dimensionalized parameters.

|  |  |  |  |
| --- | --- | --- | --- |
| $F_{\text{probe}}$ | 25–200 | [N] | Varied in steps of 25 units. |
| $R_{\text{farfield}}$ | 2.5 | $\mu\text{m}$ | Chosen based on initial test simulations. |
| $\nu$ | 0.5 | unitless | Poisson ratio of soft biological tissue [Lin et al., 2007]. |
| $R_{\text{probe}}$ | 2.5 | $\mu\text{m}$ | Based on values in Van Helvert and Friedl [2016]. |
| $k_{\text{pillar}}$ | 1 | $[\text{N m}^{-1}]$ | Chosen based on parameter scan. |

## Data analysis

During simulation, the cellular Potts spin configuration and the molecular dynamics fiber particle positions were exported at regular intervals, allowing to fully reconstruct the simulation state at the selected timepoints. For all analyses and graphical representation of simulation data, we used Python 3.8.10 and the Python libraries pandas, scikit-image, scikit-learn, numpy, and matplotlib to restructure, quantify, and plot the data [Harris et al., 2020, Hunter, 2007, McKinney, 2010, The Pandas Development Team, 2020, Van der Walt et al., 2014, Van Rossum and Drake Jr, 1995]. Quantification graphics were saved as PDF files and assembled into multi-panel figures using Inkscape v.1.1.2. When using data from external sources, we performed minor aesthetic adjustments without changing the underlying data.

### Graphical representation of simulation state

To graphically represent the simulation state, we plotted segments between subsequent fiber particles as lines; crosslinkers were plotted as lines connecting fiber particles involved in the crosslinker bond; adhesion-associated beads were plotted as circles. The cellular Potts configuration was overlayed as a semi-transparent layer using filled contour plots. For the plots of colored fibers on the right in Fig. 2 B, each fiber was assigned a unique identifier and color based on its position in the simulated domain at the initial time-point. Plots from the same simulation at later timepoints were generated by recovering the fiber identifier and its unique color assignment.

### Spatial map of fiber density

For each fiber segment, we generated 100 points between subsequent fiber particles via interpolation of the particles’ position. All fiber particles’ positions and the interpolated points were then used to generate a 2D histogram with bins subdividing the spatial domain. Spatial bins with a higher number of points thus represented a higher fiber density. Note that this approach is equivalent to experimentally measuring fluorescent markers incorporated at regular intervals into the fiber monomers.

### Densification factor

We first defined two areas on a spatial grid over the simulated domain: First, a concentric region near the cell was chosen such that it ranged 1.25 μm inside, and 5 μm outside the cell boundary. This was done by repeatedly applying morphological operators of erosion or dilation on the cellular Potts configuration. Second, an annular region far from the cell was chosen to have an outer radius tangent to the spatial domain boundary, and an inner radius 5 μm smaller. The areas defined in this way were used to generate binary array masks which we multiplied with the spatial map of fiber density generated as described in the previous section. To obtain the average density, we simply divided the sum of the masked spatial map of fiber density by the respective area. Finally, for each timepoint, the densification factor *f*_dens_ was calculated as the average fiber density in the region concentric to the cell’s boundary ⟨*ρ*_near_⟩ divided by the average fiber density in the annular region at the simulation domain’s boundary ⟨*ρ*_far_⟩:

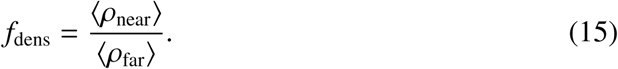

### Fiber displacement

For each fiber particle *i* we denoted its initial position as *x*_*i*_(*t* = *t*_0_) and its position at time *t* as *x*_*i*_(*t*). Then the net displacement at time *t* for that particle was

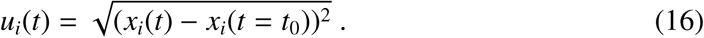

To generate quiverplots (Fig. 2 B, Fig. 4 B, Supplementary Fig. 4, 5, 6), we calculated each fiber particle’s net displacement; the quiver arrows were scaled down to reduce plot crowding while maintaining relative arrow length proportions. For the data in Fig. 4 C, we averaged *u*_*i*_(*t*) over several simulations.

### Displacement decay plots

We calculated the net displacement at *t* = 8 h for each fiber particle as in Equation 16. For each simulation, we obtained concentric radial bins around the cell’s boundary based on the final cell lattice occupancy configuration. Each bin had a width of 6.25 μm. Then, we assigned each fiber particle to one of these bins based on its final position after rounding to get integer coordinates. The average net displacement of all particles *i* contained in a bin *k* was calculated as

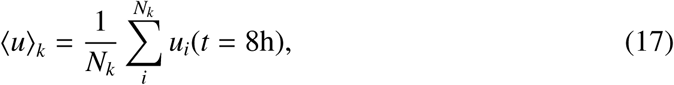

where the sum was over the number of fiber particles *N*_*k*_ in bin *k*. To normalize the data, we divided by the maximum value of all bins in a particular simulation

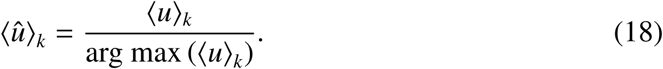

Finally, for the data in Fig. 4 E, we plotted the average and standard deviation of ⟨*û*⟩_*k*_ based on several replicate simulations.

### In silico AFM sector statistics

We subdivided the cell’s surroundings into sectors of equal angle *θ* relative to the horizontal axis crossing the cell centroid (Fig. 5 C, scheme on the right). For each sector *θ*_*i*_, we computed the mean Young’s modulus ⟨*E* (*θ*_*i*_)⟩ at *t* = 4 h per sector *θ*_*i*_ ≤ *θ* < *θ*_*i*+1_ within a distance *r* < *r*_cutoff_. We chose *r*_cutoff_ = 5 μm. For each simulation run, we defined ‘stiffened’ sectors as sectors where ⟨*E* (*θ*_*i*_)⟩(*r* < *r*_cutoff_) was above the 75% quantile of all measured sectors in that run. Other sectors were defined as ‘unstiffened’. We background-corrected these data by subtracting the mean background modulus of the network in the absence of cell-applied forces at *t* = 0 h.

## Author contributions

ET - software, simulation, analysis, figures, writing, funding; BHB - conceptualization, software, simulation, analysis, figures, writing; KAEK - software, writing; HJH - conceptualization, supervision, funding; RMHM - conceptualization, supervision, writing, funding

## Acknowledgements

This work was supported by NWO grant NWO/ENW-VICI 865.17.004 to RMHM (ET, KK, and RMHM). ET has received support from Leiden University Fund ‘Subsidy wetenschappelijk project’ grant number W213078-1, www.luf.nl. HJH acknowledges support from the Netherlands Organization for Scientific Research (NWO) (grant 639.032.612). Initial work for this manuscript was carried out on the Dutch national e-infrastructure with the support of SURF Cooperative. The remainder of this work was performed using the compute resources from the Academic Leiden Interdisciplinary Cluster Environment (ALICE) provided by Leiden University.

## Supplementary Data

**Supplementary Figure 1:**
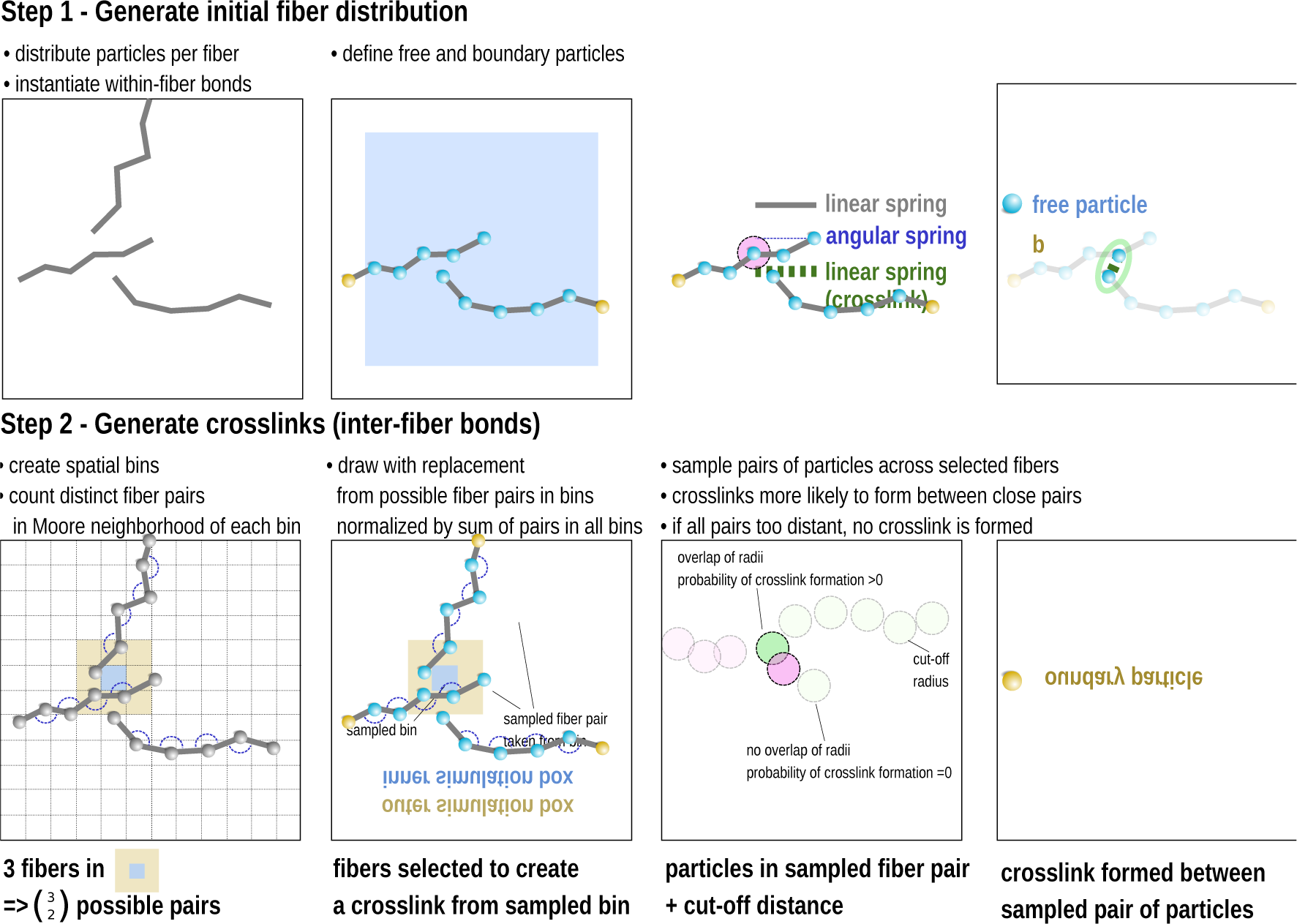
Network construction rules. We initialize the network in two steps, as depicted. After initialization and before adding the CPM model interface, we allow the network to mechanically equilibrate by integrating for *t*_init_ steps.

**Supplementary Figure 2:**
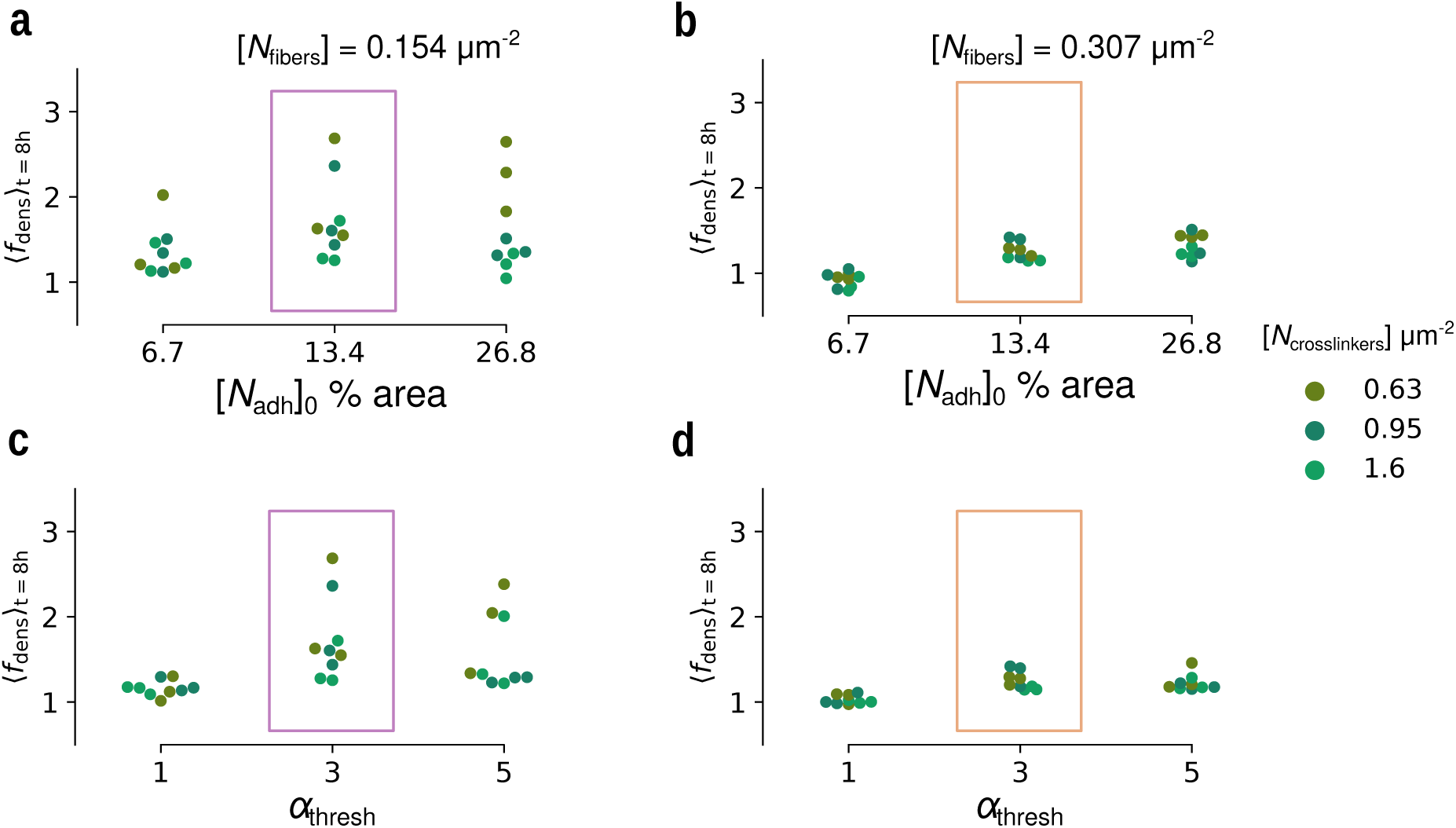
Fiber densification for several parameter choices. **a-b** Densification in final simulation state plotted against initial adhesion density for simulations with low ([*N*_fibers_] = 0.154 μm^−2^) or high ([*N*_fibers_] = 0.307 μm^−2^) fiber concentration. **c-d** Densification in final simulation state plotted against maximum penalty-free adhesion clustering for simulations with low ([*N*_fibers_] = 0.154 μm^−2^) or high ([*N*_fibers_] = 0.307 μm^−2^) fiber concentration. Highlights indicate repeated datasets in a and c; and in b and d.

**Supplementary Figure 3:**
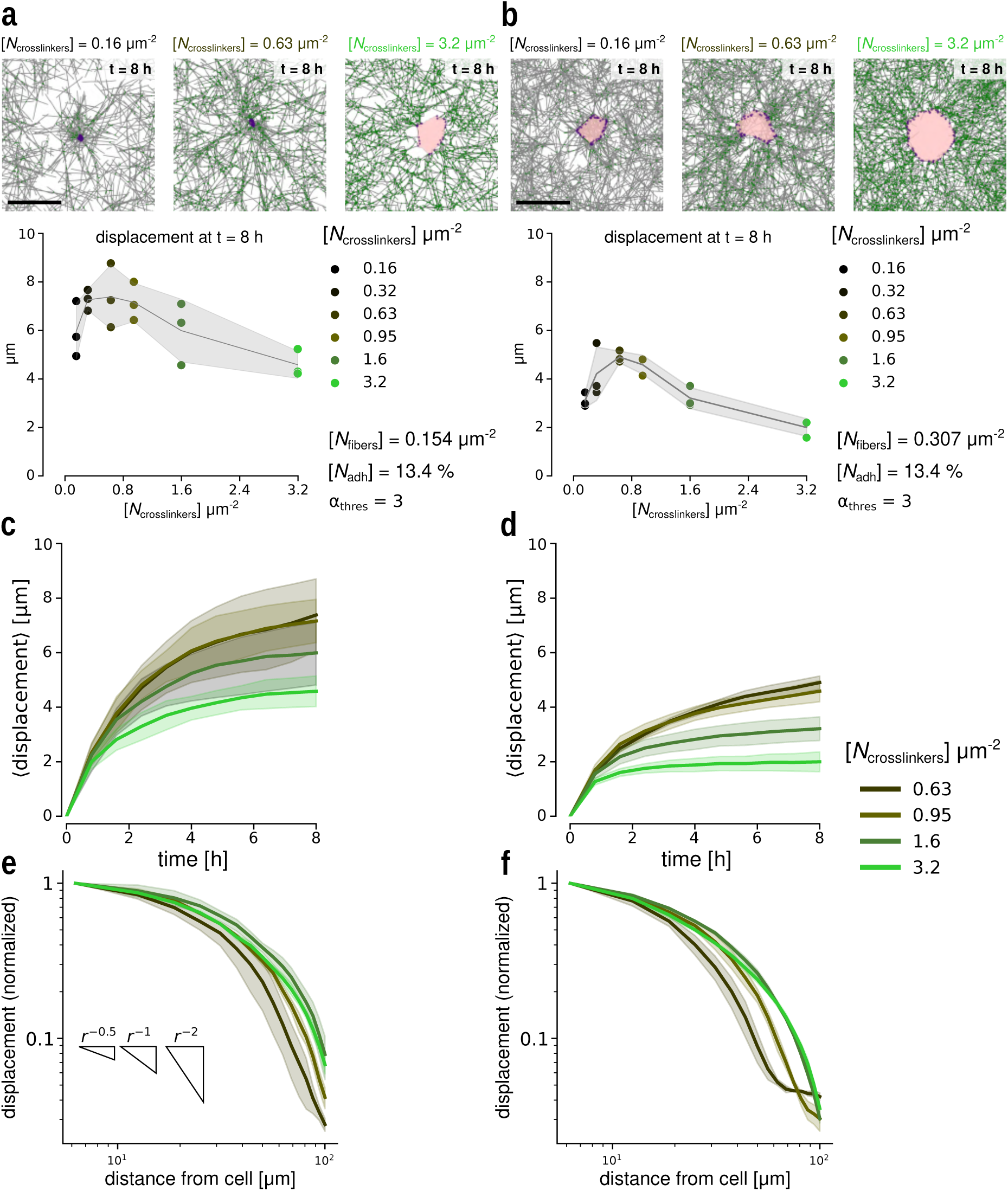
Fiber displacement at low and high fiber concentration. **a-b** Top: Final simulation state for contractile cells in relatively sparse or dense networks (respectively [*N*_fibers_] = 0.154 μm^−2^ or 0.307 μm^−2^). Bottom: Total displacement averaged in a 30 μm radius from the cell’s center at the final simulation state plotted against crosslinker density; mean and standard deviation of three simulations for each point. **c-d** Average displacement over time at low and high fiber density (respectively [*N*_fibers_] = 0.154 μm^−2^ or 0.307 μm^−2^); mean and standard deviation of three simulations each. **e-f** Total displacement averaged in 25 μm bins at the final simulation state plotted against distance from the cell edge at low and high fiber density (respectively [*N*_fibers_] = 0.154 μm^−2^ or 0.307 μm^−2^; mean and standard deviation of three simulations each. Scale bars: 20 μm.

**Supplementary Figure 4:**
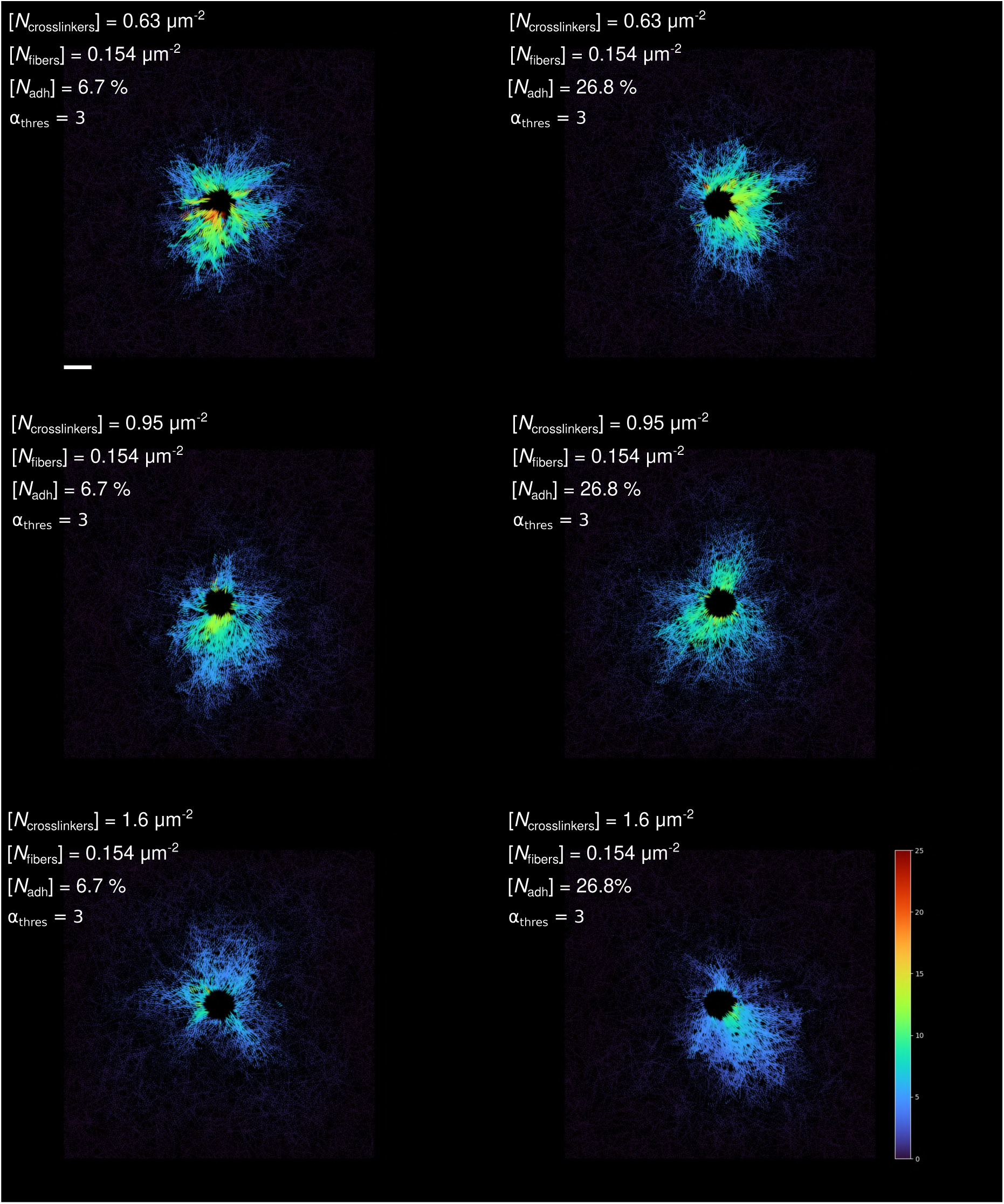
Fiber displacement at low fiber concentration. Quiverplots showing total displacement of individual fiber particles at the final simulation state with respect to the initial state. Columns show increasing initial adhesion concentration, rows have increasing crosslinker density. Simulation parameters: [*N*_fibers_] = 0.154 μm^−2^, [*α*_thresh_] = 3. Units of colorbar: μm. Scale bar: 20 μm.

**Supplementary Figure 5:**
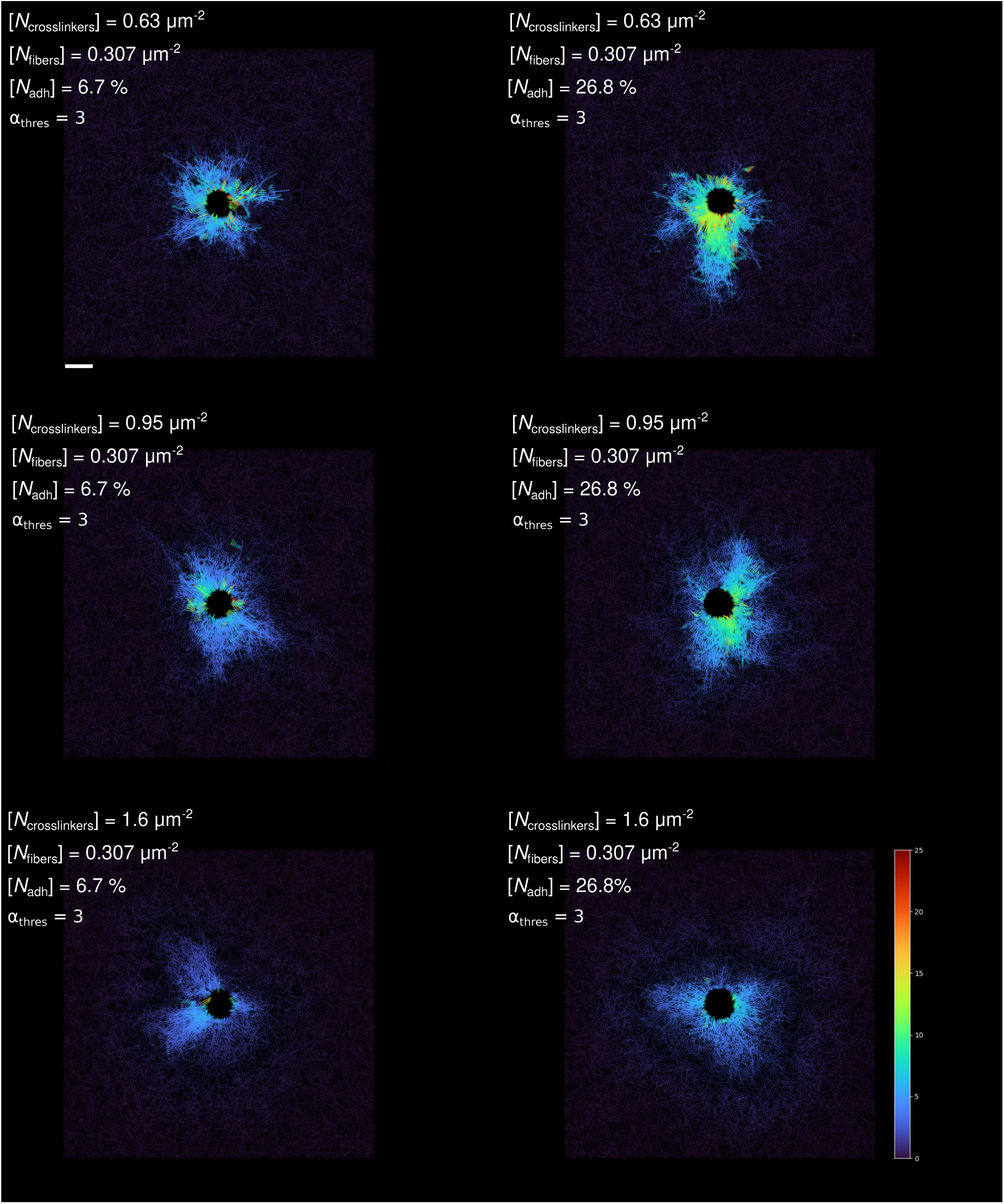
Fiber displacement at high fiber concentration. Quiverplots showing total displacement of individual fiber particles at the final simulation state with respect to the initial state. Columns show increasing initial adhesion concentration, rows have increasing crosslinker density. Simulation parameters: [*N*_fibers_] = 0.307 μm^−2^, [*α*_thresh_] = 3. Units of colorbar: μm. Scale bar: 20 μm.

**Supplementary Figure 6:**
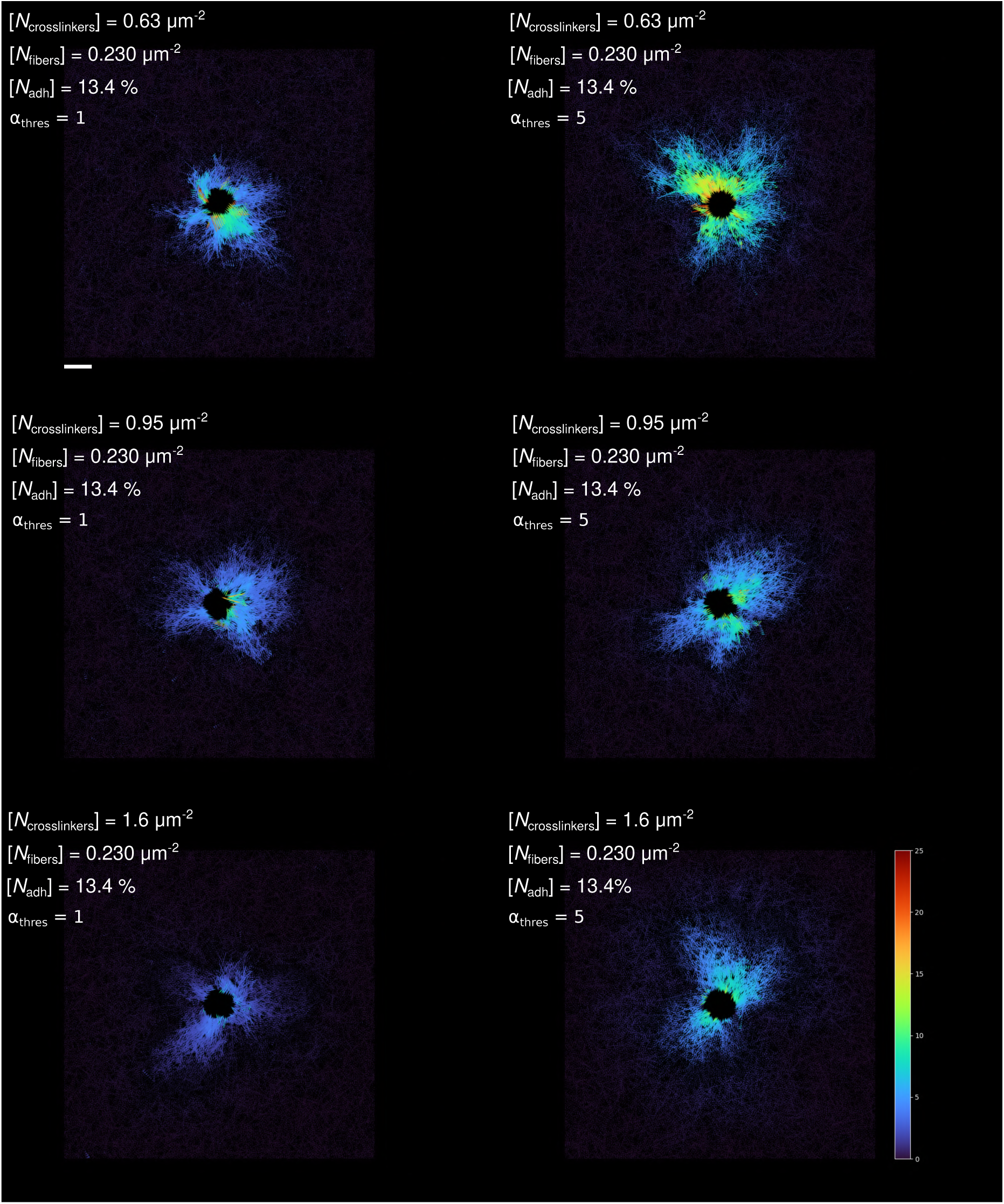
Fiber displacement at varying adhesion clustering threshold. Quiverplots showing total displacement of individual fiber particles at the final simulation state with respect to the initial state. Columns show increasing adhesion clustering capacity, rows have increasing crosslinker density. Simulation parameters: [*N*_fibers_] = 0.307 μm^−2^, [*N*_adh_]_0_ = 13.4%. Units of colorbar: μm. Scale bar: 20 μm.

**Supplementary Figure 7:**
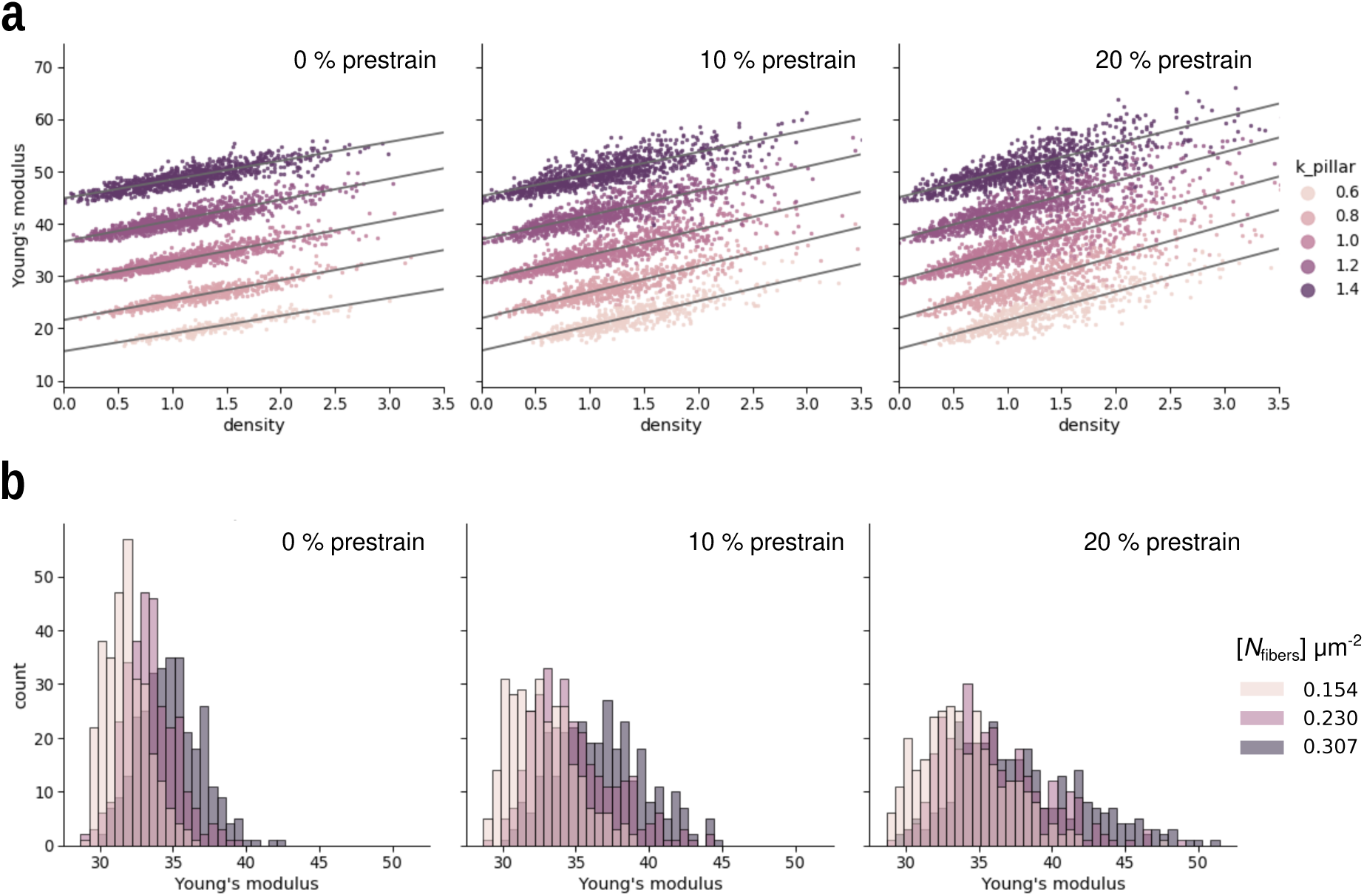
Calibration of in silico AFM. **a**. Measured Young’s modulus for varying prestrain values, as a function of local ECM density. Note the linear dependence on pillar grid elasticity *k*_pillar_. **b** Distribution of Young’s moduli in cell-free isotropic networks, with *k*_pillar_ = 1.

**Supplementary Figure 8:**
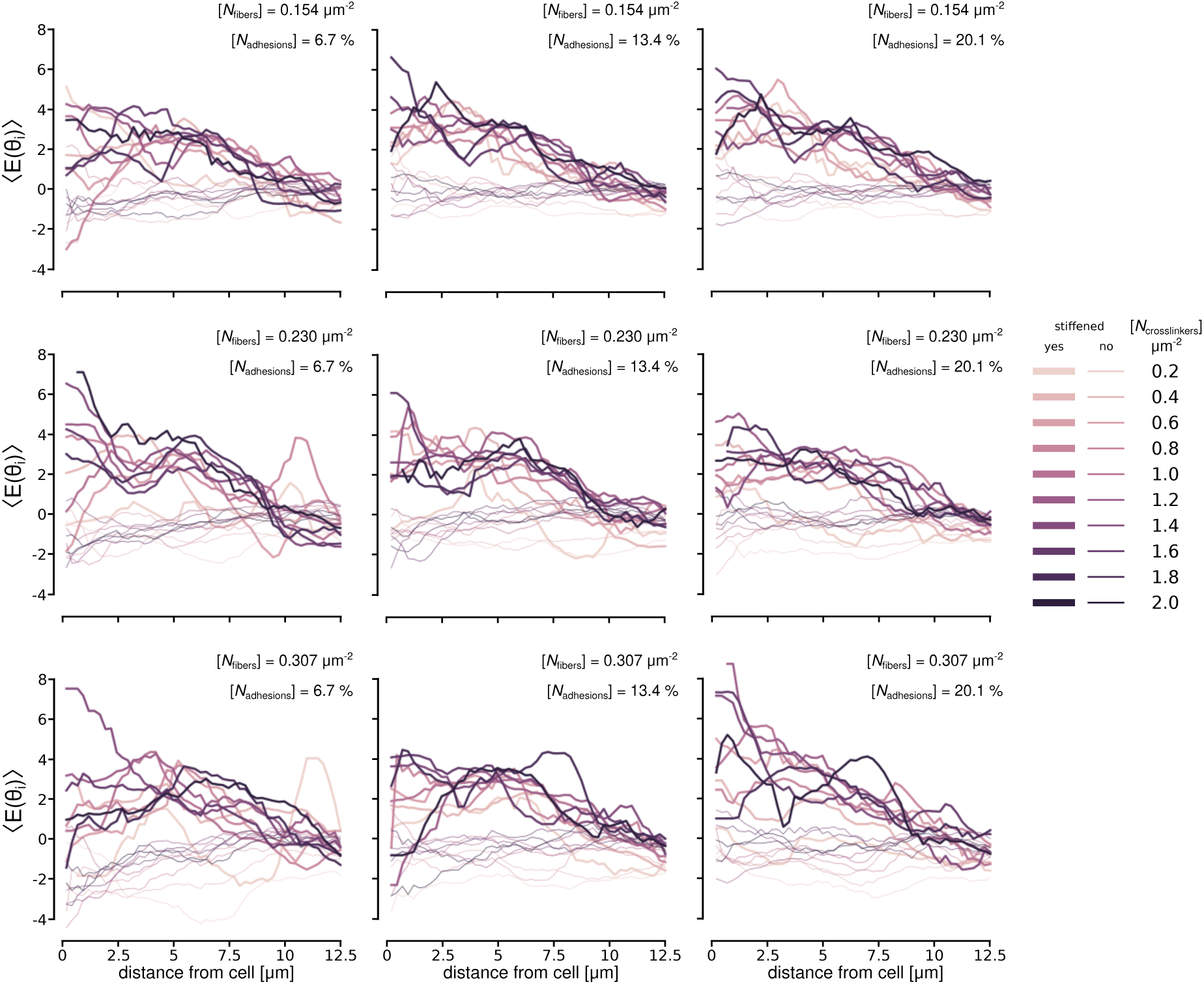
Cell-induced strain-stiffening. Young’s moduli as function of distance from the cell. Stiffened areas are defined as those above the 75% quantile of mean Young’s moduli within 5 μm from the cell perimeter. As a control the unstiffened areas are plotted and observed to agree with the background modulus. Rows correspond to increasing fiber density. Columns correspond to increasing cell-ECM adhesions. All plots use *α*_thresh_ = 3.

Supplementary Movie 1: Cell contraction at intermediate crosslinker density.

At intermediate crosslinker density, cell contraction and fiber densification is reduced. The fiber network behaves as a soft gel. Simulation corresponds to data in Figure 2 C. The movie spans 10000 timesteps (≈ 8 h) with one frame every 200 timesteps. Simulation parameters: [*N*_crosslinkers_] = 0.95 μm^−2^, *α*_thresh_ = 3, [*N*_fibers_] = 0.23 μm^−2^, [*N*_adh_]_0_ = 13.4%.

Supplementary Movie 2: Cell contraction at low crosslinker density.

At low crosslinker density, the cell contracts strongly, concentrating fibers in its immediate proximity. Simulation corresponds to data in Figure 2 C. The movie spans 10000 timesteps (≈ 8 h) with one frame every 500 timesteps. Simulation parameters: [*N*_crosslinkers_] = 0.32 μm^−2^, *α*_thresh_ = 3, [*N*_fibers_] = 0.23 μm^−2^, [*N*_adh_]_0_ = 13.4%.

Supplementary Movie 3: Cell contraction at high crosslinker density.

At high crosslinker density, the network behaves as a stiff gel preventing any substantial remodelling. Simulation corresponds to data in Figure 2 C. The movie spans 10000 timesteps (≈ 8 h) with one frame every 500 timesteps. Simulation parameters: [*N*_crosslinkers_] = 3.2 μm^−2^, *α*_thresh_ = 3, [*N*_fibers_] = 0.23 μm^−2^, [*N*_adh_]_0_ = 13.4%.

Supplementary Movie 4: Cell contraction at low fiber density.

At low fiber density, the network behaves as a softer gel. Simulation corresponds to data in Figure 3 C. The movie spans 10000 timesteps (≈ 8 h) with one frame every 200 timesteps. Simulation parameters: [*N*_crosslinkers_] = 0.95 μm^−2^, *α*_thresh_ = 3, [*N*_fibers_] = 0.154 μm^−2^, [*N*_adh_]_0_ = 13.4%.

Supplementary Movie 5: Cell contraction at high fiber density.

At low fiber density, the network behaves as a stiffer gel. Simulation corresponds to data in Figure 3 C. The movie spans 10000 timesteps (≈ 8 h) with one frame every 200 timesteps. Simulation parameters: [*N*_crosslinkers_] = 0.95 μm^−2^, *α*_thresh_ = 3, [*N*_fibers_] = 0.307 μm^−2^, [*N*_adh_]_0_ = 13.4%.

Supplementary Movie 6: Cell contraction with fewer initial cell-ECM adhesions.

Cell contraction is reduced with fewer initial adhesions. Simulation corresponds to data in Figure 3 D. The movie spans 10000 timesteps (≈ 8 h) with one frame every 500 timesteps. Simulation parameters: [*N*_crosslinkers_] = 0.95 μm^−2^, *α*_thresh_ = 3, [*N*_fibers_] = 0.230 μm^−2^, [*N*_adh_]_0_ = 6.7%.

Supplementary Movie 7: Cell contraction with more initial cell-ECM adhesions.

Cell contraction is not appreciably increased past a certain amount of initial adhesions. Simulation corresponds to data in Figure 3 D. The movie spans 10000 timesteps (≈ 8 h) with one frame every 500 timesteps. Simulation parameters: [*N*_crosslinkers_] = 0.95 μm^−2^, *α*_thresh_ = 3, [*N*_fibers_] = 0.230 μm^−2^, [*N*_adh_]_0_ = 26.8%.

Supplementary Movie 8: Cell contraction with less adhesion clustering.

Cell contraction is reduced if the adhesion clustering threshold is lowered. Simulation corresponds to data in Figure 3 E. The movie spans 10000 timesteps (≈ 8 h) with one frame every 500 timesteps. Simulation parameters: [*N*_crosslinkers_] = 0.95 μm^−2^, *α*_thresh_ = 1, [*N*_fibers_] = 0.230 μm^−2^, [*N*_adh_]_0_ = 13.4%.

Supplementary Movie 9: Cell contraction with more adhesion clustering.

Cell contraction is increased if the adhesion clustering threshold is increased. Simulation corresponds to data in Figure 3 E. The movie spans 10000 timesteps (≈ 8 h) with one frame every 500 timesteps. Simulation parameters: [*N*_crosslinkers_] = 0.95 μm^−2^, *α*_thresh_ = 5, [*N*_fibers_] = 0.230 μm^−2^, [*N*_adh_]_0_ = 13.4%.

Supplementary Movie 10: Contraction-mediated fiber displacement at low crosslinker density.

At low crosslinker density, only fibers immediately proximal to the cell are affected by contraction. Simulation corresponds to data in Figure 4 A-B. The movie spans 10000 timesteps (≈ 8 h) with one frame every 500 timesteps. Simulation parameters: [*N*_crosslinkers_] = 0.16 μm^−2^, *α*_thresh_ = 3, [*N*_fibers_] = 0.230 μm^−2^, [*N*_adh_]_0_ = 13.4%.

Supplementary Movie 11: Contraction-mediated fiber displacement at intermediate crosslinker density.

At intermediate crosslinker density, visible fiber displacement extends several cell radii away from the contracting cell. Simulation corresponds to data in Figure 4 A-B. The movie spans 10000 timesteps (≈ 8 h) with one frame every 500 timesteps. Simulation parameters: [*N*_crosslinkers_] = 0.63 μm^−2^, *α*_thresh_ = 3, [*N*_fibers_] = 0.230 μm^−2^, [*N*_adh_]_0_ = 13.4%.

Supplementary Movie 12: Contraction-mediated fiber displacement at high crosslinker density.

At high crosslinker density fiber displacement spreads far, but the magnitude of the displacement is small. Simulation corresponds to data in Figure 4 A-B. The movie spans 10000 timesteps (≈ 8 h) with one frame every 500 timesteps. Simulation parameters: [*N*_crosslinkers_] = 3.2 μm^−2^, *α*_thresh_ = 3, [*N*_fibers_] = 0.230 μm^−2^, [*N*_adh_]_0_ = 13.4%.

Supplementary Movie 13: Contracting cell doublet.

Fibers between contracting cells form straightened bundles. Simulation parameters: [*N*_crosslinkers_] = 0.63μm^−2^, *α*_thresh_ = 3, [*N*_fibers_] = 0.230 μm^−2^, [*N*_adh_]_0_ = 13.4%.

## Notes

### Competing Interest Statement

The authors have declared no competing interest.

### Summary of Updates

Minor revisions to strengthen relevance statement of the methodology.

## References

P. J. Albert and U. S. Schwarz. Dynamics of cell shape and forces on micropatterned substrates predicted by a cellular potts model. Biophysical Journal, 106(11):2340–2352, 2014.

F. Alisafaei, X. Chen, T. Leahy, P. A. Janmey, and V. B. Shenoy. Long-range mechanical signaling in biological systems. Soft Matter, 17(2):241–253, 2021.

J. A. Anderson, C. D. Lorenz, and A. Travesset. General purpose molecular dynamics simulations fully implemented on graphics processing units. 227(10):5342–5359, 2008.

J. A. Anderson, J. Glaser, and S. C. Glotzer. HOOMD-blue: A python package for high-performance molecular dynamics and hard particle monte carlo simulations. Computational Materials Science, 173: 109363, 2020.

A. L. Bauer, T. L. Jackson, and Y. Jiang. A cell-based model exhibiting branching and anastomosis during tumor-induced angiogenesis. Biophysical Journal, 92(9):3105–3121, 2007.

P. Buske, J. Przybilla, M. Loeffler, N. Sachs, T. Sato, H. Clevers, and J. Galle. On the biomechanics of stem cell niche formation in the gut–modelling growing organoids. The FEBS Journal, 279(18):3475–3487, 2012.

O. Chaudhuri, J. Cooper-White, P. A. Janmey, D. J. Mooney, and V. B. Shenoy. Effects of extracellular matrix viscoelasticity on cellular behaviour. Nature, 584(7822):535–546, 2020.

E. S. Colizzi, R. M. A. Vroomans, and R. M. H. Merks. Evolution of multicellularity by collective integration of spatial information. eLife, 9:e56349, 2020.

J. T. Daub and R. M. H. Merks. A cell-based model of extracellular-matrix-guided endothelial cell migration during angiogenesis. Bulletin of Mathematical Biology, 75(8):1377–1399, 2013.

J. T. Daub and R. M. H. Merks. Cell-based computational modeling of vascular morphogenesis using tissue simulation toolkit. In Vascular Morphogenesis, pages 67–127. Springer, 2014.

C. D. Davidson, W. Y. Wang, I. Zaimi, D. K. P. Jayco, and B. M. Baker. Cell force-mediated matrix reorganization underlies multicellular network assembly. Scientific Reports, 9(1):1–13, 2019.

D. Drasdo and S. Hoehme. Modeling the impact of granular embedding media, and pulling versus pushing cells on growing cell clones. New Journal of Physics, 14(5):055025, 2012.

S.-J. Dunn, P. L. Appleton, S. A. Nelson, I. S. Näthke, D. J. Gavaghan, and J. M. Osborne. A two-dimensional model of the colonic crypt accounting for the role of the basement membrane and pericryptal fibroblast sheath. PLOS Computational Biology, 8(5):e1002515, 2012.

B. J. Dzamba and D. W. DeSimone. Extracellular matrix (ecm) and the sculpting of embryonic tissues. In Current Topics in Developmental Biology, volume 130, pages 245–274. Elsevier, 2018.

J. F. Eichinger, M. J. Grill, I. D. Kermani, R. C. Aydin, W. A. Wall, J. D. Humphrey, and C. J. Cyron. A computational framework for modeling cell–matrix interactions in soft biological tissues. Biomechanics and Modeling in Mechanobiology, 20(5):1851–1870, 2021.

L. Feld, L. Kellerman, A. Mukherjee, A. Livne, E. Bouchbinder, and H. Wolfenson. Cellular contractile forces are nonmechanosensitive. Science Advances, 6(17):eaaz6997, 2020.

J. A. Glazier and F. Graner. Simulation of the differential adhesion driven rearrangement of biological cells. Physical Review E, 47(3):2128, 1993.

F. Graner and J. A. Glazier. Simulation of biological cell sorting using a two-dimensional extended potts model. Physical Review Letters, 69(13):2013, 1992.

Y. Guo, S. Calve, and A. B. Tepole. Multiscale mechanobiology: Coupling models of adhesion kinetics and nonlinear tissue mechanics. Biophysical Journal, 121(4):525–539, 2022.

Y. L. Han, P. Ronceray, G. Xu, A. Malandrino, R. D. Kamm, M. Lenz, C. P. Broedersz, and M. Guo. Cell contraction induces long-ranged stress stiffening in the extracellular matrix. Proceedings of the National Academy of Sciences, 115(16):4075–4080, 2018.

A. K. Harris, D. Stopak, and P. Wild. Fibroblast traction as a mechanism for collagen morphogenesis. Nature, 290(5803):249–251, 1981.

C. R. Harris, K. J. Millman, S. J. Van der Walt, R. Gommers, P. Virtanen, D. Cournapeau, E. Wieser, J. Taylor, S. Berg, N. J. Smith, et al. Array programming with numpy. Nature, 585(7825):357–362, 2020.

S. D. Hester, J. M. Belmonte, J. S. Gens, S. G. Clendenon, and J. A. Glazier. A multi-cell, multi-scale model of vertebrate segmentation and somite formation. PLOS Computational Biology, 2011.

T. Hirashima, E. G. Rens, and R. M. H. Merks. Cellular potts modeling of complex multicellular behaviors in tissue morphogenesis. Development, Growth & Differentiation, 59(5):329–339, 2017.

U. Horzum, B. Ozdil, and D. Pesen-Okvur. Step-by-step quantitative analysis of focal adhesions. MethodsX, 1:56–59, 2014.

D. L. Humphries, J. A. Grogan, and E. A. Gaffney. Mechanical cell–cell communication in fibrous networks: the importance of network geometry. Bulletin of Mathematical Biology, 79(3):498–524, 2017.

J. D. Hunter. Matplotlib: A 2d graphics environment. Computing in Science & Engineering, 9(3):90–95, 2007. doi: 10.1109/MCSE.2007.55.

T. Korff and H. G. Augustin. Tensional forces in fibrillar extracellular matrices control directional capillary sprouting. Journal of Cell Science, 112(19):3249–3258, 1999.

S. T. Kreger, B. J. Bell, J. Bailey, E. Stites, J. Kuske, B. Waisner, and S. L. Voytik-Harbin. Polymerization and matrix physical properties as important design considerations for soluble collagen formulations. Biopolymers: Original Research on Biomolecules, 93(8):690–707, 2010.

K. A. Leonidakis, P. Bhattacharya, J. Patterson, B. E. Vos, G. H. Koenderink, J. Vermant, D. Lambrechts, M. Roeffaers, and H. Van Oosterwyck. Fibrin structural and diffusional analysis suggests that fibers are permeable to solute transport. Acta Biomaterialia, 47:25–39, 2017.

D. C. Lin, E. K. Dimitriadis, and F. Horkay. Robust strategies for automated afm force curve analysis—i. non-adhesive indentation of soft, inhomogeneous materials. Journal of Biomechanical Engineering, 129 (3):430–440, 2007.

C. K. Macnamara, A. Caiazzo, I. Ramis-Conde, and M. A. J. Chaplain. Computational modelling and simulation of cancer growth and migration within a 3d heterogeneous tissue: The effects of fibre and vascular structure. Journal of Computational Science, 40:101067, 2020.

A. Malandrino, X. Trepat, R. D. Kamm, and M. Mak. Dynamic filopodial forces induce accumulation, damage, and plastic remodeling of 3d extracellular matrices. PLOS Computational Biology, 15(4): e1006684, 2019.

A. Mann, R. S. Sopher, S. Goren, O. Shelah, O. Tchaicheeyan, and A. Lesman. Force chains in cell–cell mechanical communication. Journal of the Royal Society Interface, 16(159):20190348, 2019.

A. F. M. Marée, A. Jilkine, A. Dawes, V. A. Grieneisen, and L. Edelstein-Keshet. Polarization and movement of keratocytes: a multiscale modelling approach. Bulletin of Mathematical Biology, 68(5):1169–1211, 2006.

M. Marin-Riera, M. Brun-Usan, R. Zimm, T. Välikangas, and I. Salazar-Ciudad. Computational modeling of development by epithelia, mesenchyme and their interactions: a unified model. Bioinformatics, 32 (2):219–225, 2016.

D. E. Mason, J. M. Collins, J. H. Dawahare, T. D. Nguyen, Y. Lin, S. L. Voytik-Harbin, P. Zorlutuna, M. C. Yoder, and J. D. Boerckel. Yap and taz limit cytoskeletal and focal adhesion maturation to enable persistent cell motility. Journal of Cell Biology, 218(4):1369–1389, 2019.

W. McKinney. Data Structures for Statistical Computing in Python. In Stéfan Van der Walt and Jarrod Millman, editors, Proceedings of the 9th Python in Science Conference, pages 56–61, 2010.

R. M. H. Merks and J. A. Glazier. A cell-centered approach to developmental biology. Physica A, 2005.

J. Metzcar, Y. Wang, R. Heiland, and P. Macklin. A review of cell-based computational modeling in cancer biology. JCO Clinical Cancer Informatics, 2:1–13, 2019.

M. Michael and M. Parsons. New perspectives on integrin-dependent adhesions. Current Opinion in Cell Biology, 63:31–37, 2020.

S. R. Naganathan, M. Popović, and A. C. Oates. Left–right symmetry of zebrafish embryos requires somite surface tension. Nature, pages 1–6, 2022.

B. K. A. Nelemans, M. Schmitz, H. Tahir, R. M. H. Merks, and T. H. Smit. Somite Division and New Boundary Formation by Mechanical Strain. iScience, 2020.

T. J. Newman. Modeling multicellular systems using subcellular elements. Mathematical Biosciences And Engineering, 2(3):613–624, 00 2005.

I. Niculescu, J. Textor, and R. J. De Boer. Crawling and gliding: a computational model for shape-driven cell migration. PLOS Computational Biology, 11(10):e1004280, 2015.

J. Notbohm, A. Lesman, P. Rosakis, D. A. Tirrell, and G. Ravichandran. Microbuckling of fibrin provides a mechanism for cell mechanosensing. Journal of The Royal Society Interface, 12(108):20150320, 2015.

K. H. Palmquist, S. F. Tiemann, F. L. Ezzeddine, S. Yang, C. R. Pfeifer, A. Erzberger, A. R. Rodrigues, and A. E. Shyer. Reciprocal cell-ecm dynamics generate supracellular fluidity underlying spontaneous follicle patterning. Cell, 2022.

D. Panja. Anomalous polymer dynamics is non-markovian: memory effects and the generalized langevin equation formulation. Journal of Statistical Mechanics: Theory and Experiment, 2010(06):P06011, 2010.

D. Peurichard, F. Delebecque, A. Lorsignol, C. Barreau, J. Rouquette, X. Descombes, L. Casteilla, and P. Degond. Simple mechanical cues could explain adipose tissue morphology. Journal of Theoretical Biology, 429:61–81, 2017.

J. W. Reinhardt and K. J. Gooch. Agent-based modeling traction force mediated compaction of cell-populated collagen gels using physically realistic fibril mechanics. Journal of Biomechanical Engineering, 136(2):021024, 2014.

J. W. Reinhardt and K. J. Gooch. An agent-based discrete collagen fiber network model of dynamic traction force-induced remodeling. Journal of Biomechanical Engineering, 140(5), 2018.

E. G. Rens and L. Edelstein-Keshet. From energy to cellular forces in the cellular potts model: An algorithmic approach. PLOS Computational Biology, 15(12):e1007459, 2019.

E. G. Rens and R. M. H. Merks. Cell contractility facilitates alignment of cells and tissues to static uniaxial stretch. Biophysical Journal, 112(4):755–766, 2017.

E. G. Rens and R. M. H. Merks. Cell shape and durotaxis explained from cell-extracellular matrix forces and focal adhesion dynamics. iScience, 23(9):101488, 2020.

E. G. Rens, M. T. Zeegers, I. Rabbers, A. Szabó, and R. M. H. Merks. Autocrine inhibition of cell motility can drive epithelial branching morphogenesis in the absence of growth. Philosophical Transactions of the Royal Society B, 375(1807):20190386, 2020.

P. Ronceray, C. P. Broedersz, and M. Lenz. Fiber networks amplify active stress. Proceedings of the National Academy of Sciences, 113(11):2827–2832, 2016.

M. S. Rudnicki, H. A. Cirka, M. Aghvami, E. A. Sander, Q. Wen, and K. L. Billiar. Nonlinear strain stiffening is not sufficient to explain how far cells can feel on fibrous protein gels. Biophysical Journal, 105(1):11–20, 2013.

S. A. Sandersius and T. J. Newman. Modeling cell rheology with the Subcellular Element Model. Physical Biology, 2008.

N. Sasaki and S. Odajima. Stress-strain curve and young’s modulus of a collagen molecule as determined by the x-ray diffraction technique. Journal of Biomechanics, 29(5):655–658, 1996.

N. J. Savill and P. Hogeweg. Modelling morphogenesis: From single cells to crawling slugs. Journal of Theoretical Biology, 184(3):229–235, 00 1997.

K. Schakenraad, G. I. Martorana, B. H. Bakker, L. Giomi, and R. M. H. Merks. Stress fibers orient traction forces on micropatterns: A hybrid cellular potts model study. bioRxiv, 2022.

D. K. Schlüter, I. Ramis-Conde, and M. A. J. Chaplain. Computational modeling of single-cell migration: the leading role of extracellular matrix fibers. Biophysical Journal, 103(6):1141–1151, 2012.

M. Scianna, L. Preziosi, and K. Wolf. A cellular potts model simulating cell migration on and in matrix environments. Mathematical Biosciences & Engineering, 10(1):235, 2013.

L. E. Scott, L. A. Griggs, V. Narayanan, D. E. Conway, C. A. Lemmon, and S. H. Weinberg. A hybrid model of intercellular tension and cell–matrix mechanical interactions in a multicellular geometry. Biomechanics and Modeling in Mechanobiology, pages 1–17, 2020.

V. D. Sree and A. B. Tepole. Computational systems mechanobiology of growth and remodeling: integration of tissue mechanics and cell regulatory network dynamics. Current Opinion in Biomedical Engineering, 15:75–80, 2020.

J. Steinwachs, C. Metzner, K. Skodzek, N. Lang, I. Thievessen, C. Mark, S. Münster, K. E. Aifantis, and B. Fabry. Three-dimensional force microscopy of cells in biopolymer networks. Nature Methods, 13(2): 171–176, 2016.

T. Sütterlin, E. Tsingos, J. Bensaci, G. N. Stamatas, and N. Grabe. A 3d self-organizing multicellular epidermis model of barrier formation and hydration with realistic cell morphology based on episim. Scientific Reports, 7:43472, 2017.

R. B. Svensson, T. Hassenkam, P. Hansen, and S. P. Magnusson. Viscoelastic behavior of discrete human collagen fibrils. Journal of the Mechanical Behavior of Biomedical Materials, 3(1):112–115, 2010.

O. Tange. GNU Parallel 2018. Ole Tange, Mar. 2018. ISBN 9781387509881.

The Pandas Development Team. pandas-dev/pandas: Pandas. Zenodo, Feb. 2020.

S. Van der Walt, J. L. Schönberger, J. Nunez-Iglesias, F. Boulogne, J. D. Warner, N. Yager, E. Gouillart, and T. Yu. scikit-image: image processing in python. PeerJ, 2:e453, 2014.

S. Van Helvert and P. Friedl. Strain stiffening of fibrillar collagen during individual and collective cell migration identified by afm nanoindentation. ACS Applied Materials & Interfaces, 8(34):21946–21955, 2016.

S. Van Helvert, C. Storm, and P. Friedl. Mechanoreciprocity in cell migration. Nature Cell Biology, 20(1): 8–20, 2018.

R. F. M. Van Oers, E. G. Rens, D. J. LaValley, C. A. Reinhart-King, and R. M. H. Merks. Mechanical cellmatrix feedback explains pairwise and collective endothelial cell behavior in vitro. PLOS Computational Biology, 10(8):e1003774, 2014.

G. Van Rossum and F. L. Drake Jr. Python Tutorial. Centrum voor Wiskunde en Informatica Amsterdam, The Netherlands, 1995.

L. Van Steijn, I. M. N. Wortel, C. Sire, L. Dupré, G. Theraulaz, and R. M. H. Merks. Computational modelling of cell motility modes emerging from cell-matrix adhesion dynamics. PLoS Computational Biology, 2022. ISSN 1553-734X.

D. A. C. Walma and K. M. Yamada. The extracellular matrix in development. Development, 147(10), 2020.

H. Wang and X. Xu. Continuum elastic models for force transmission in biopolymer gels. Soft Matter, 16 (48):10781–10808, 2020.

I. M. N. Wortel, I. Niculescu, P. M. Kolijn, N. S. Gov, R. J. de Boer, and J. Textor. Local actin dynamics couple speed and persistence in a cellular potts model of cell migration. Biophysical Journal, 120(13): 2609–2622, 2021.

